# How to barcode (almost all) freshwater biodiversity

**DOI:** 10.1101/2023.12.13.571596

**Authors:** Orianne Tournayre, Haolun Tian, David R. Lougheed, Matthew J.S. Windle, Sheldon Lambert, Jennipher Carter, Zhengxin Sun, Jeff Ridal, Yuxiang Wang, Brian F. Cumming, Shelley E. Arnott, Stephen C. Lougheed

## Abstract

Freshwater ecosystems are complex, diverse and face a variety of imminent threats that have led to changes in both ecosystem structure and function. It is urgent that we develop and standardize monitoring tools allowing for rapid and comprehensive assessment of freshwater communities to understand their changing dynamics and to inform conservation. Environmental DNA surveys offer a means to inventory and monitor aquatic diversity, yet most studies focus on one or a few taxonomic groups only. In this study, we sought to 1) identify thoroughly validated, cost-efficient primer pair combinations that maximize detection of broad swaths of freshwater diversity, and 2) facilitate future primer pair selection by creating a free online and user-friendly tool. We first evaluated the completeness of public reference sequence databases and the efficiency of 14 primer pairs using an *in silico* approach, and then performed eDNA surveys using five mock communities (mix of DNA from tissues), water samples from aquarium samples with known taxonomic composition, and finally water samples from freshwater systems in Eastern Canada. We highlight the power of eDNA-based metabarcoding for reconstructing freshwater communities, including prey, parasite, pathogen, invasive, and declining species. Our work reveals the importance of the marker choice on species resolution, as well as the importance of degenerate primers, the length of the target fragment and the filtering parameters on detection success in water eDNA samples. Our new online tool SNIPe revealed that 13 to 14 primer pairs are necessary to recover 100% of the species in water samples (aquarium and natural systems), but four primer pairs are sufficient to recover almost 75% of taxa with little overlap. These results highlight the usefulness of eDNA for freshwater monitoring and should prompt more studies on tools to survey all-inclusive communities.

## INTRODUCTION

Despite covering less than 1% of the Earth’s surface, freshwater ecosystems play a disproportionate role in biogeochemical and ecological processes (Dudgeon et al. 2006). They are incredibly diverse comprising more than 400 large-scale ecoregions (Abell et al. 2008), rivers, lakes, and wetlands being home to more than 10% of all described species on Earth (fungi, plants, invertebrates, and vertebrates; Reid et al. 2019; Albert et al. 2021). Freshwater ecosystems also provide critical services to humans including drinking water, food, transportation, sanitation, and hydroelectric energy. The heavy reliance of human societies on freshwater has been a key driver of freshwater ecosystem decline over the past century: habitat destruction, pollution, introduction of invasive species, overharvesting, all overlain by climate change, have contributed to rapid declines of freshwater biodiversity (IPBES, 2019). For example, 70% of the world’s wetlands have been lost since 1900 (Davidson, 2014) and one third of the freshwater species assessed by the International Union for Conservation of Nature (IUCN) Red List are threatened with extinction (IUCN, 2017). There is no one-fits-all solution to protect freshwater ecosystems because of their complexity, diversity, high levels of endemism, and heterogeneity of threats they face, varying in frequency, magnitude and duration (Ahmed et al. 2022; Dudgeon et al. 2006). These perturbations can markedly disrupt ecological structure and function (Angeler et al. 2014; Dudgeon et al. 2006), so questions pertaining to biodiversity conservation and stewardship of aquatic ecosystems necessarily involve quantification of interactions among species (e.g. effect of invasive species on native communities, altered trophic complexity and trophic cascades; Higgins and Zanden 2010; Karatayev and Burlakova 2022; Piczak et al. 2023). It is crucial that we develop, validate and standardize monitoring tools that allows rapid and comprehensive assessment of freshwater communities to better characterize community composition and change in responses to perturbations to inform conservation (Ficetola and Taberlet 2022).

Environmental DNA (eDNA) based molecular approaches are increasingly used as tools for freshwater monitoring complementing traditional approaches that rely on collecting organisms like seining (Andres et al. 2023; Berger et al. 2020; Gibson et al. 2023), benthic invertebrate surveys (Duarte et al. 2021; Ji et al. 2022), or electrofishing (Goutte et al. 2020). While current eDNA-based approaches are limited to presence/absence or estimates of relative abundance (Beng and Corlett 2020; Zaiko et al. 2018), they typically capture a greater range of species, including those that are rare and elusive, or detect some species not captured by traditional approaches (Fediajevaite et al. 2021; Hallam et al. 2021; Keck et al. 2021; Lyet et al. 2021; Sales et al. 2020). The growing importance of eDNA methods to support freshwater conservation is highlighted by the fact that national and international conservation organizations and companies have begun to develop eDNA-based programs or services. For example, the IUCN has launched *eBioAtlas*, a program that aims to infill global freshwater biodiversity gaps using cutting-edge eDNA technology on >30,000 freshwater samples from around the world. However, in almost 20 years of eDNA studies, evaluations of entire biotic communities remain rare, probably because of the substantial technical challenges and methodological limitations (Ficetola and Taberlet 2022). The two major methodological limitations that influence detection success (species presence/absence) and the taxonomic resolution of such eDNA surveys are: i) incomplete reference databases, and ii) limits to taxonomic specificity and complementarity of the primer pair(s). Reference databases are usually geographically and taxonomically biased (Marques et al. 2021; Weigand et al. 2019) and this can only be solved by more collaborations between taxonomists and molecular biologists, such as The Barcode of Life initiative aiming at creating a public collection of reference sequences from vouchered specimens of all species of life (Costa and Carvalho 2007; Ratnasingham and Hebert 2007). Databases must also be carefully curated and quality-checked as user-entered data can contain many errors associated with the production and archival of the sequences (Keck, Couton, and Altermatt 2023). There is no consensus on the best strategy to overcome primer bias such as PCR affinity (Freeland 2017). Current approaches combine group-specific primer pairs, universal primer pairs, or both universal and group-specific primer pairs, on one or several genes (e.g. COI and 12S) (see review in Ficetola and Taberlet 2022). While increasing the number of primer pairs increases the number of detected taxa, it also increases the cost, the labour, and time of analyses, meaning all-inclusive community studies are rarely applied in monitoring programs. Ultimately, there is a trade-off between the number of primers to combine and the number of taxa detected.

Our case study focuses on the St. Lawrence River area in Ontario, Canada. The St. Lawrence river drains the Laurentian Great Lakes which contain 21% of the world’s surface freshwater; a system that has been affected by diverse anthropogenic stressors that have had widespread impacts on its environmental health including biodiversity (Smith et al. 2019). Lost of riparian habitats and coastal wetlands, as well as habitat degradation, are particularly acute in the area of study with a loss of approximately 70% of historic wetlands in South Ontario (Marbek 2023). We had two objectives: 1) to identify suites of cost-efficient primer pairs that maximize our ability to quantify species richness within freshwater water bodies, spanning plankton, macrophytes, invertebrates and vertebrates, and 2) create a free online tool to optimize primer pair selection for future surveys using either published or new eDNA detection data as we obtained for Objective 1.

## MATERIAL AND METHODS

### 1. In silico evaluation

#### 1.1. Selection of the genes and primer pairs: completeness of reference databases

We identified the gene(s) with the most exhaustive reference database(s) for Eastern Canadian vertebrates (reptiles, amphibians, mammals, birds, fish), invertebrates (zooplankton, annelids and molluscs), aquatic plants and phytoplankton (e.g. cyanobacteria) by comparing the number of sequences available in NCBI (GenBank; Benson et al. 2008) and BOLD (Ratnasingham and Hebert 2007) first in 2021, and then updated in 2023 before our final analyses. Six genes were evaluated: mitochondrial cytochrome oxidase 1 – COI (GenBank, BOLD), mitochondrial cytochrome b – cytB (GenBank), mitochondrial 12S rRNA (GenBank), mitochondrial 16S rRNA (GenBank), nuclear 18S rRNA (GenBank) and chloroplast ribulose bisphosphate carboxylase rbcl (GenBank). Insects were not included in this analysis as the COI gene has already been identified as the ‘gold standard’ for DNA barcoding (Elbrecht et al. 2019; Hebert, Ratnasingham, and de Waard 2003; Meusnier et al. 2008; Porter and Hajibabaei 2018; Ratnasingham and Hebert 2007; Yu et al. 2012).

#### 1.2. In silico evaluation of the sequences

We downloaded all GenBank and BOLD sequences of zooplankton, annelid, platyhelminth, mollusc, reptile, amphibian, fish, mammal, and bird families occurring in Ontario and Quebec using PrimerMiner (Elbrecht and Leese 2016). The sequences of 13 orders of freshwater invertebrates, previously used in Elbrecht and Leese (2016, 2017) were also downloaded: Plecoptera, Ephemeroptera, Trichoptera, Coleoptera, Odonata, Megaloptera, Amphipoda, Isopoda, Diptera, Turbellaria, Polychaeta, Oligochaeta and Hirudinea. As the program could not accommodate all sequence data simultaneously, a threshold of 2000 downloaded sequences was used as recommended by Elbrecht and Leese (2016). When available, the mitochondrial reads were aligned separately (i.e., for each group of taxa) using Multiple Alignment using Fast Fourier Transform (MAFFT) as implemented in Geneious Prime v7.1.9 (Kearse et al. 2012), and the OTUs were mapped to the mitochondrial consensus sequence. The visualization of the alignments was made using PrimerMiner. The numbers of OTUs per group of taxa are detailed in Table S1.

We selected group-specific and universal primer pairs from the literature (Table 1): i) three universal mitochondrial COI primer pairs (Folmer et al. 1994; Leray et al. 2013; Tournayre et al. 2020), ii) one 16S rRNA (Klymus et al. 2017) and two COI (Elbrecht and Leese 2017; Leese et al. 2021) primer pairs for macroinvertebrates, iii) one 12S rRNA (Miya et al. 2015) and one COI (Roy et al. 2018) fish primer pair, iv) one rbcl plant primer pair (Coghlan, Shafer, and Freeland 2021), and v) one 18S rRNA (Zhan et al. 2013) and one bacterial 16S rRNA (Armingohar et al. 2014) primer pair for plankton (cyanobacteria, diatoms, dinoflagellates, ciliates, rotifers, and crustaceans). Based on the sequence alignments, we modified four of these primer pairs to increase their *in silico* amplification success (e.g., degenerate bases to prevent mismatches): “16SMOL2”, “COIFishdegen”, “Uniorbcl” and “UniPh” (Table 1).

**Table 1.**
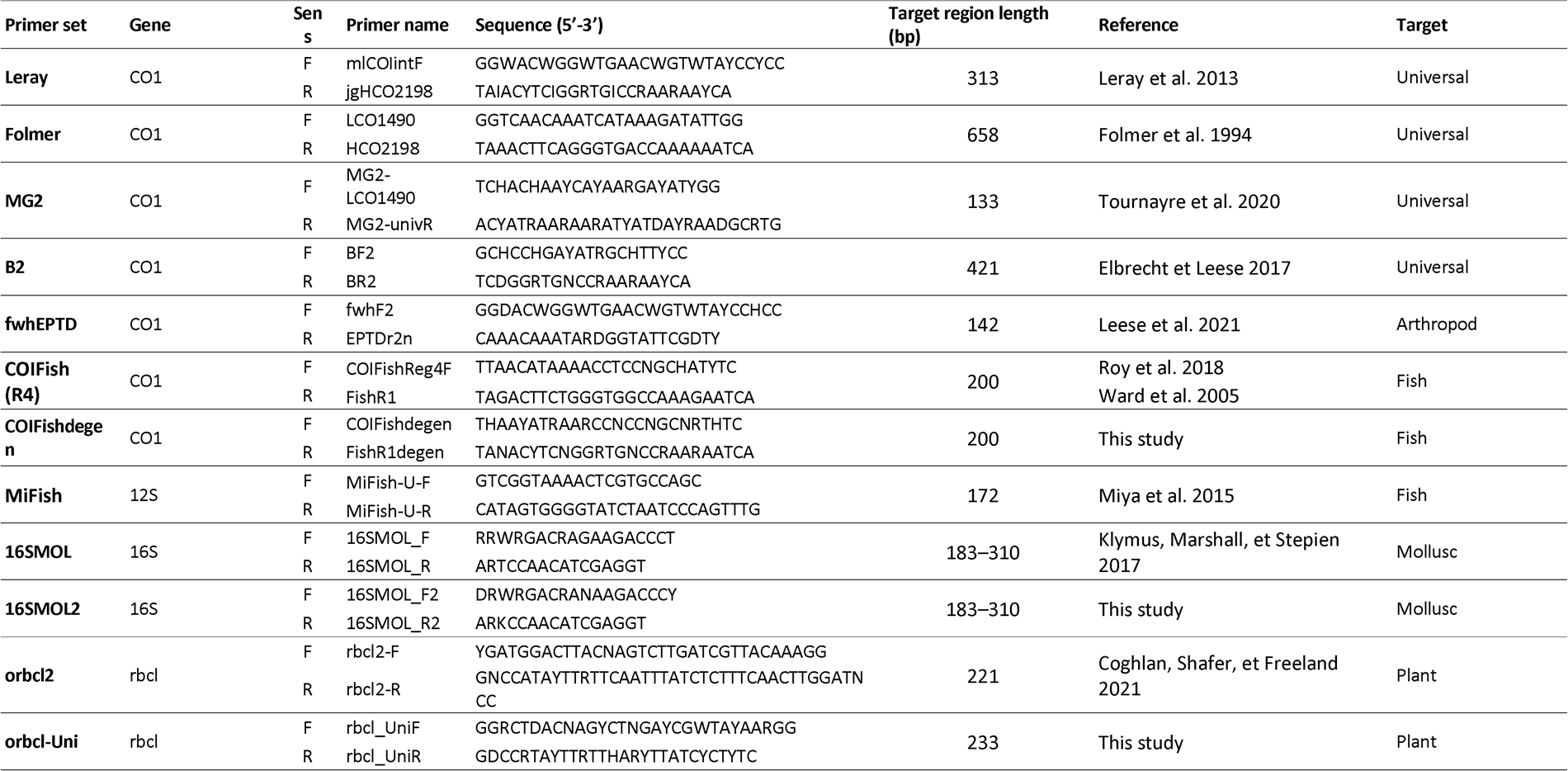
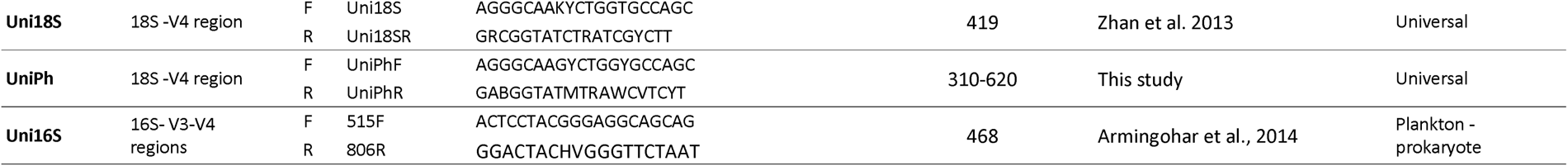
Details of primer pairs including primer names, target regions, sequences, focal taxa, amplicon size, and references.

Forward and reverse primer evaluations were done individually for each taxon using PrimerMiner. Unlike other software, PrimerMiner considers the position of mismatches (with higher penalties for 3’ mismatches), the type of mismatch (e.g., G–A and A–G have more severe impacts on PCR amplification; Stadhouders et al. 2010) and whether two mismatches are adjacent (adjacent mismatches lead to higher instability). The higher the penalty score calculated using PrimerMiner, the worse the primer is expected to perform. The mean penalty scores (MPS) of both primers were summed per taxon (forward + reverse).

### 2. Mock community preparation and sequencing

#### 2.1. Sample collection and DNA extraction

Tissue and DNA samples were provided by the Royal Ontario Museum (ethanol preserved tissues), the River Institute (frozen tissues from the Fish Identification Nearshore Survey program), Thousand Island National Park (frozen tissue from road kills), the Canadian Phycological Culture Centre (specimen from cultures), and from Queen’s University (frozen and ethanol-preserved tissues, and DNA stored at -80°C). Detailed information on samples (type of samples, year, storage, and DNA extraction methods) is available in Table S2.

#### 2.2. Composition of the mock communities (MC)

DNA concentration of each sample was assessed using the DeNovix dsDNA High Sensitivity 2 Points Assay kit, following manufacturer protocols. We created five mock communities (MC) using 145 taxa (see Table S3 for MC composition) : i) MC_1_ included all taxa with a concentration ≥ 5ng/µL (N = 91), ii) MC_2_ included only vertebrates with a concentration ≥ 5ng/µL (N = 73), iii) MC_3_ included only invertebrates with a concentration ≥ 5ng/µL (N = 8), iv) MC_4_ included only plants and plankton with a concentration ≥ 5ng/µL (N = 9), and v) MC_5_ included all tissue-derived DNA samples with a concentration ≤ 5ng/µL (N = 61). One mammal species (gray four-eyed opossum, *Philander opossum*) was included in MC_1_ and MC_2_, and was used as an alien control to estimate the false assignment rate (see section 5. Bioinformatics and taxonomic assignments).

### 3. eDNA water samples

We collected 1L water samples from seven tanks at the Brockville Aquatarium (Ontario, Canada) on October 22^th^, 2021 using sterile 1L Nalgene bottles. Disposable nitrile gloves were worn during sampling and changed between tanks. All equipment was decontaminated before and after use in a 10% bleach bath overnight and rinsed thoroughly with distilled water. The first tank contained ten fish species (“BA1”, water volume = 20,000gal, Table 2). The water system of the second tank (“BA2”, one species only: Longnose gar, *Lepisosteus osseus*) was shared with three other tanks (“Shoreline”, “Lake”, “Sturgeon”) for a total of 20 fish species and a water volume of 40,000 gallons (Table 2). The third tank (“BA4”) contained Sea lamprey (*Petromyzon marinus*). The fourth tank (“BA5”, Eastern musk turtle, *Sternotherus odoratus*) shared a water system with the grey tree frog (*Dryophytes versicolor*) tank. The fifth tank (“BA6”, eastern garter snake, *Thamnophis sirtalis sirtalis*) shared its water system with the American eel (*Anguilla rostrata*) tank. The sixth tank had American bullfrog (“BA7”, *Lithobates catesbeianus*). The seventh tank (“BA8”, Spotted turtle *Clemmys guttata*) shared the water system with three other tanks (Leopard frog, *Lithobates pipiens*, Green frog, *Lithobates clamitans*, and Mink frog, *Lithobates septentrionalis*). Water samples were stored in a cooler with ice packs for return to the laboratory (under four hours) for filtration.

**Table 2.**
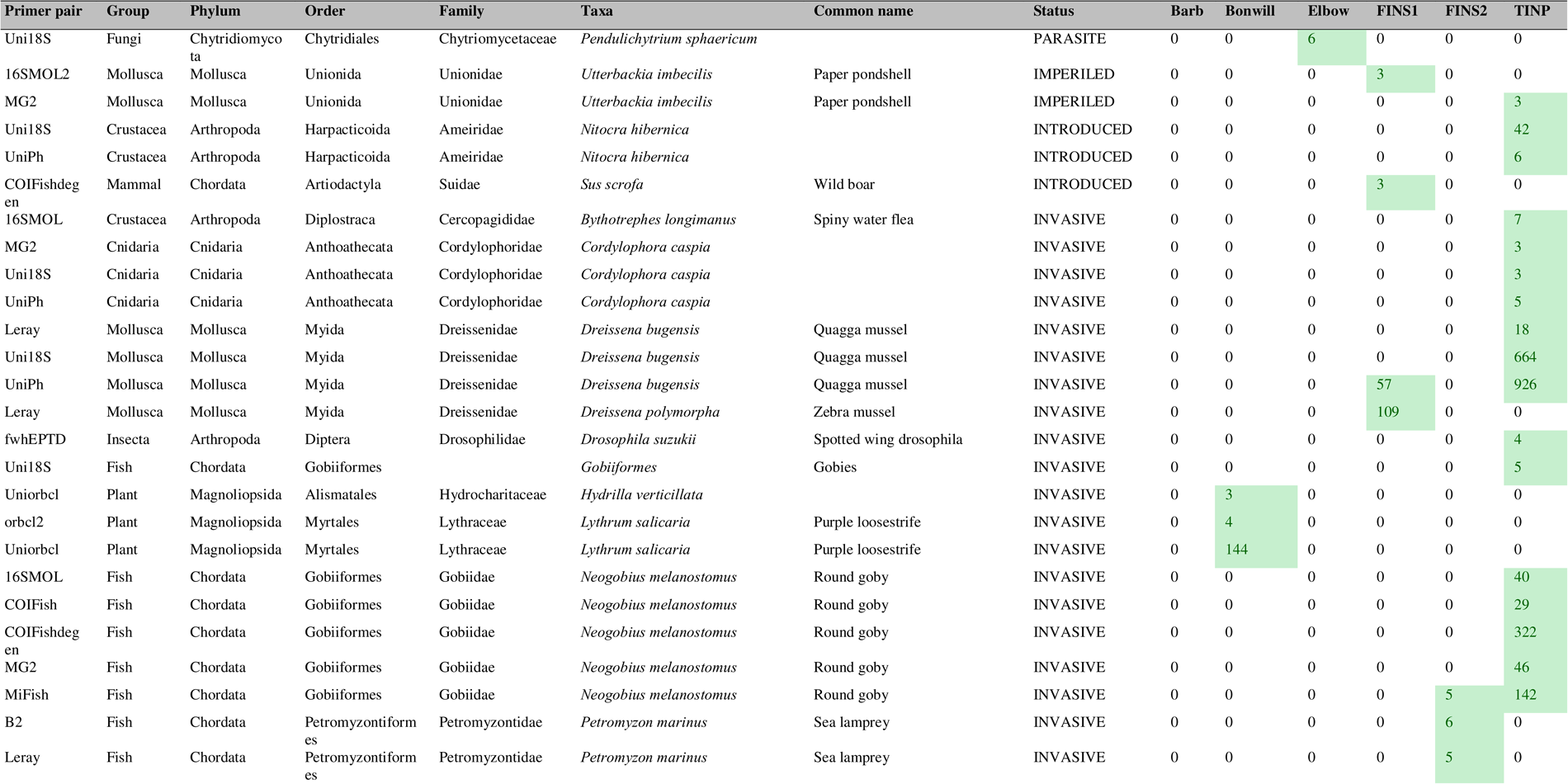
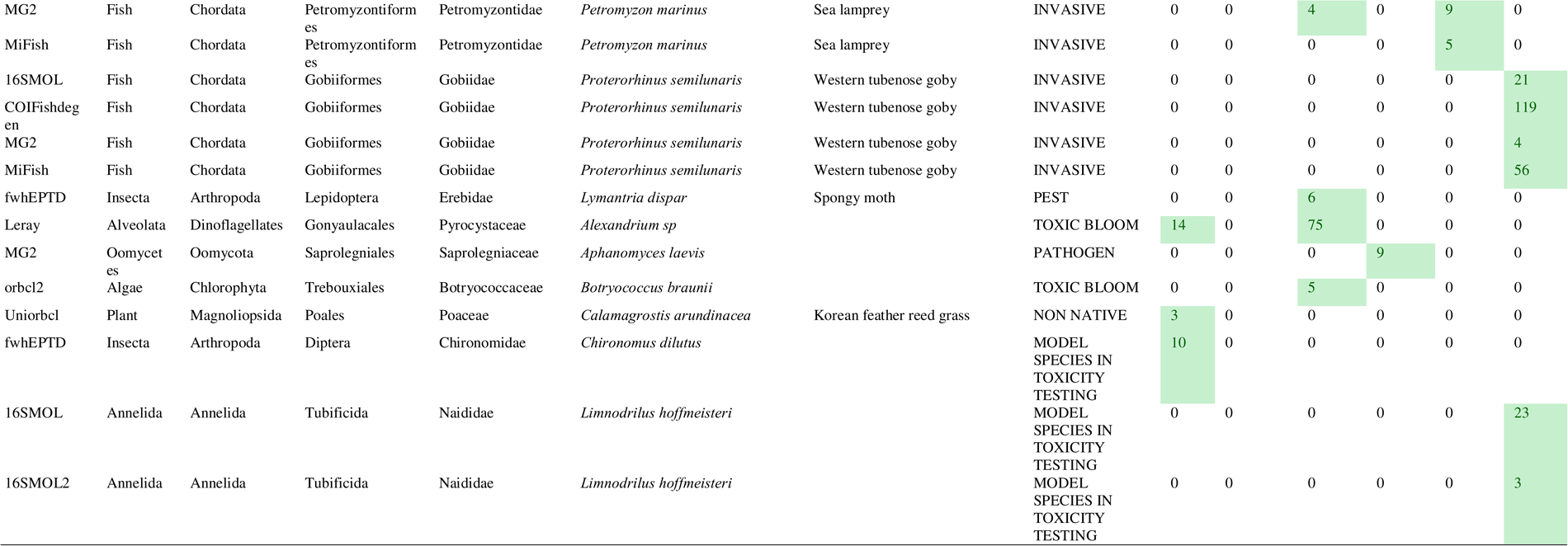
Number of reads of the taxa of interest relevant to conservation, human health, and agriculture, at the six sites and for each primer pair. “FINS1” = Pointes des cascades, “FINS2” = Hoople NW, “TINP” = Thousand Islands National Park.

Finally, we collected three 1L surface water samples from six waterbodies in southeastern Ontario (Canada) varying in type, size and landscape context to maximise the detection of a broad range of species: one upland marsh “Barb” (2021/06/02, 44.52526/-76.37179), one woodland pond “Bonwill” (2021/06/03, 44.542148/-76.399359), one small lake “Elbow” (2021/06/06, 44.473328/-76.430148) plus three sites on the St Lawrence River : “FINS1” (Pointes-des-cascades, 2020/08/06, 45.33436462/-73.95526161), “FINS2” (Hoople NW, 2020/09/04, 44.98663186/-74.94213128) and “TINP” (Thousand Island National Park, 2021/10/27, 44.45343717/-75.85865145). The equipment (including waders) was bleached (10%) and rinsed thoroughly with distilled water before and after sampling between sites. Disposable nitrile gloves were worn for all sampling. The samples were stored in a cooler with ice packs for return to the lab for the filtration (< six hours).

We filtered water using polycarbonate filter membranes (PCTE; pore size: 1.0 μm, Sterilitech, Auburn, WA, USA) housed in a 47mm in-line filter holder (Pall, Port Washington, NY, USA) with Masterflex 24 tubing (Masterflex, Gelsenkirchen, Germany) and a peristaltic pump (Wattera, Missisauga, ON, Canada). Prior to sample filtration and between filtration events, 1L of 10% bleach followed by 1L of distilled water were processed to decontaminate the filter holder and tubing (i.e., to avoid cross-contamination between samples). The filter membranes were folded with sterile tweezers (dipped in 95% ethanol and flamed twice to disinfect) and individually stored in 2 mL microcentrifuge tubes containing 500 μL 2% (w/v) cetrimonium bromide extraction buffer (CTAB). Filters were stored at -20°C until eDNA extraction. The eDNA extractions of the filters were conducted using a chloroform-based method (Turner et al. 2014) with the following modifications: the DNA pellet was eluted in 100 µL pre-warmed TE buffer (55°C) and the 100 µL eDNA extracts were combined by site for a final volume of 300 µL. eDNA and negative controls of filtration and extraction were stored at -20 °C until library preparation.

### 4. Library preparation

We used a two-step PCR protocol to amplify target regions and index the samples (Galan et al. 2018). We included a PCR negative control for each primer pair and batch of 95 samples. PCR_1_ was performed in 20 µL reaction volumes using 10 µL of 2X Taq FroggaMix (FroggaBio, Concord, ON, Canada), 5 µL of nuclease free water (FroggaBio), 1 µL of forward primer (10µM), 1 µL of reverse primer (10µM), and 3 µL of eDNA extract. PCR_1_ cycling conditions consisted of an initial denaturation step at 95°C for 15 min, followed by 40 cycles of denaturation at 94°C for 30 s, annealing at 45°C for 45 s, and extension at 72°C for 2 min, followed by a final extension step at 72°C for 10 min (Tournayre et al. 2020). PCR_1_ products were checked on an agarose gel (1.5%) stained with RedSafe™. PCR_2_ was carried out in 24 µL reaction volumes using 12 µL of 2X Taq FroggaMix (FroggaBio, Concord, ON, Canada), 8 µL of nuclease free water, 1 µL of each indexed primers i5 (10µM) and i7 (10µM) (8 bases each; Kozich et al. 2013) and 2 µL of PCR_1_ product. PCR_2_ started with an initial denaturation step of 95°C for 15 min, followed by 10 cycles of denaturation at 95°C for 40 s, annealing at 55°C for 45 s and extension at 72°C for 2 min followed by a final extension step at 72°C for 10 min. PCR_2_ products were again checked on an agarose gel (1.5%). For each primer pair, we pooled the replicates and samples per sample type: aquarium samples, waterbody samples, and mock community samples. We loaded 60 µL of each of these 37 libraries on an agarose gel (1.0%) stained with RedSafe™ and then excised bands with single-use scalpel blades on a UV transilluminator. Gel slices were placed in 2.0 mL sterile tubes and purified using a Wizard® SV Gel and PCR Clean-Up System (Promega, Madison, WI, USA) following the manufacturer’s protocol. The 37 purified libraries were quantified using the NEBNext® Library Quant Kit for Illumina® (NEB E7630, New England Biolabs, Ipswich, MA, USA) kit on a Bio-Rad CFX96 qPCR instrument (Bio-Rad Laboratories, Hercules, CA, USA) and pooled in equimolar proportion to obtain a final library at 4 nM. 10 pM of the library and 10% of PhiX control were sequenced on a MiSeq Illumina flow cell with a 600-cycle Reagent v3 kit (2x300bp pair-end sequencing, Illumina, San Diego, CA, USA).

### 5. Bioinformatics and taxonomic assignments

Raw sequencing reads were demultiplexed using the MiSeq Reporter Software based on the index combinations. We used the v1.7.6 Barque pipeline (Mathon et al. 2021) to process the reads (settings are detailed in Appendix S1). Briefly, raw reads were trimmed using Trimmomatic v0.36 (Bolger, Lohse, and Usadel 2014), paired-end reads were merged using Flash v1.2.11 (Magoč and Salzberg 2011), and merged-reads were split based on primer pair. Sequences of unexpected length were discarded, as were chimeric sequences using Vsearch v2.15.2 (Rognes et al. 2016). Unique reads were merged and assigned to species using Vsearch (≥98% identity, ≥60% coverage). We discarded all species highly unlikely to be present in our study region, including via lab contamination (e.g., polar bear, *Ursus marinus*), marine species, species not occurring in Canada or within our study area, micro-organisms in the mock communities (e.g. *Wolbachia*, diatoms, cyanobacteria, crustaceans such as the freshwater ostracod, *Cypridopsis vidua*) that were likely unintentionally sampled along with target taxa. Due to uncertainty on species taxonomy, we considered bacterial taxa only at the genus level, excluding cyanobacteria. We then used 1) detections from the negative controls to filter for cross-contamination (T_CC;_ Galan et al. 2016), 2), the alien control (gray four-eyed opossum, *P. opossum*) to filter positive detections due to the false assignment rate of reads (T_FA;_ Galan et al. 2016), and 3) the technical replicates to control for PCR and sequencing stochasticity (we considered a positive detection for a taxon to comprise at least two positive PCR replicates out of three), and 4) we only retained taxa with at least three reads in the eDNA water samples as an additional step to decrease false positive detections while minimizing false negatives. Last, we discarded potential redundancy: when sequences were identified at both the species level and the order or family or genus level, we kept only the identification at the species level (note that in some cases this strategy discarded a high number of reads; for example 4,528 reads for a midge, *Dicrotendipes* sp. and the primer pair fwhEPTD). We kept detection at the genus level if the potential species was not already identified at the species level (e.g. 16SMOL detected *Ameiurus natalis* and *Ameiurus melas/nebulosus* so we kept both) or if the genus level identification was detected in a different sample than the species level identification (e.g. *Tanytarsus* sp. for primer pair fwhEPTD).

### 6. Statistical analyses

All the statistical analyses were done in R v4.3.1 (R Core Team 2022). We ran a quasi-binomial generalized linear regression model (GLM) and subsequent Analysis of Deviance Table to assess the effect of the number of reads, target fragment size (bp), the percentage of degenerate bases in the primer sequences (forward + reverse), and the total number of expected taxa as a proxy of the mock community complexity on the proportion of detected taxa per taxonomic group (excluding null proportion). As we did not expect any particular number of detected taxa for the water eDNA samples collected in southeastern Ontario, we used instead a quasi-Poisson GLM and the number of detected taxa as the response variable (excluding null detections).

A quasi-binomial GLM model and subsequent Analysis of Deviance Table were performed to assess the effect of the gene, target fragment size (bp) and the percentage of degenerate bases in the primer sequences (forward + reverse) on the percentage of taxa identified at the species level. Note that we considered only the expected species that passed all the filtering criteria in the mock community and aquarium analyses. We assessed the significance of differences between pairs of group means by conducting a post-hoc analysis with the *emmeans* package (Tukey method for p-value adjustment) (Lenth 2023).

### 7. Selecting Novel Informative Primer-sets for eDNA (SNIPe)

We developed a new online tool with a graphical user interface for eDNA primer pair selection based on empirical species detection data from multiple primer pairs that may vary in taxonomic focus and resolution: https://snipe.dlougheed.com/.

This new tool involves two steps: 1) “Choose or upload a dataset” allowing use of our default dataset using 14 primer pairs or by uploading new datasets or data from the literature; 2) “Discover primer sets” allowing users to choose the suite of taxa of interest and a maximum number of primer pairs desired, and then to explore the number of OTUs detected for combinations of primer pairs from one to the maximum specified. Results include the percentage of recovered taxa per primer pair combination, an indication of which taxa would be added using ‘n’ primer pairs versus combinations with ‘n-1’ primer pairs, Euler diagrams to visualize overlap among primer pairs, data on taxonomic resolution for each primer pair combination, and a stacked histogram showing taxon accumulation for different numbers of primer pairs. The default eDNA dataset (water samples from the six sites in Eastern Ontario) is available in Table S5.

## RESULTS

### 1. *In silico* evaluation: reference database and performance

#### 1.1. Gene(s) with the most exhaustive reference(s) database(s)

We assessed a total of 1031 taxa spanning 144 orders and 257 families: 27 reptiles, 26 amphibians, 180 fish, 16 semi-aquatic mammals, 72 aquatic birds, 101 molluscs, 33 annelids and platyhelminths, five rotifers, 59 crustaceans, 116 plants and 396 phytoplankton taxa (Table S6). COI gene had the most exhaustive reference database for vertebrates and invertebrates (Figure 1A). Two species of branchiopods, one malacostracans, four annelids, one mollusc and seven fish had no sequence available for any genes (Table S6). All plant species had rbcl sequences available (Figure 1B). Bacterial 16S had the most exhaustive reference database for cyanobacteria and 18S for eukaryotic phytoplankton taxa (Figure 1C). However, more than one third of the phytoplankton species (37.8%) did not have sequence available for any of the tested genes (Table S6).

**Figure 1.**
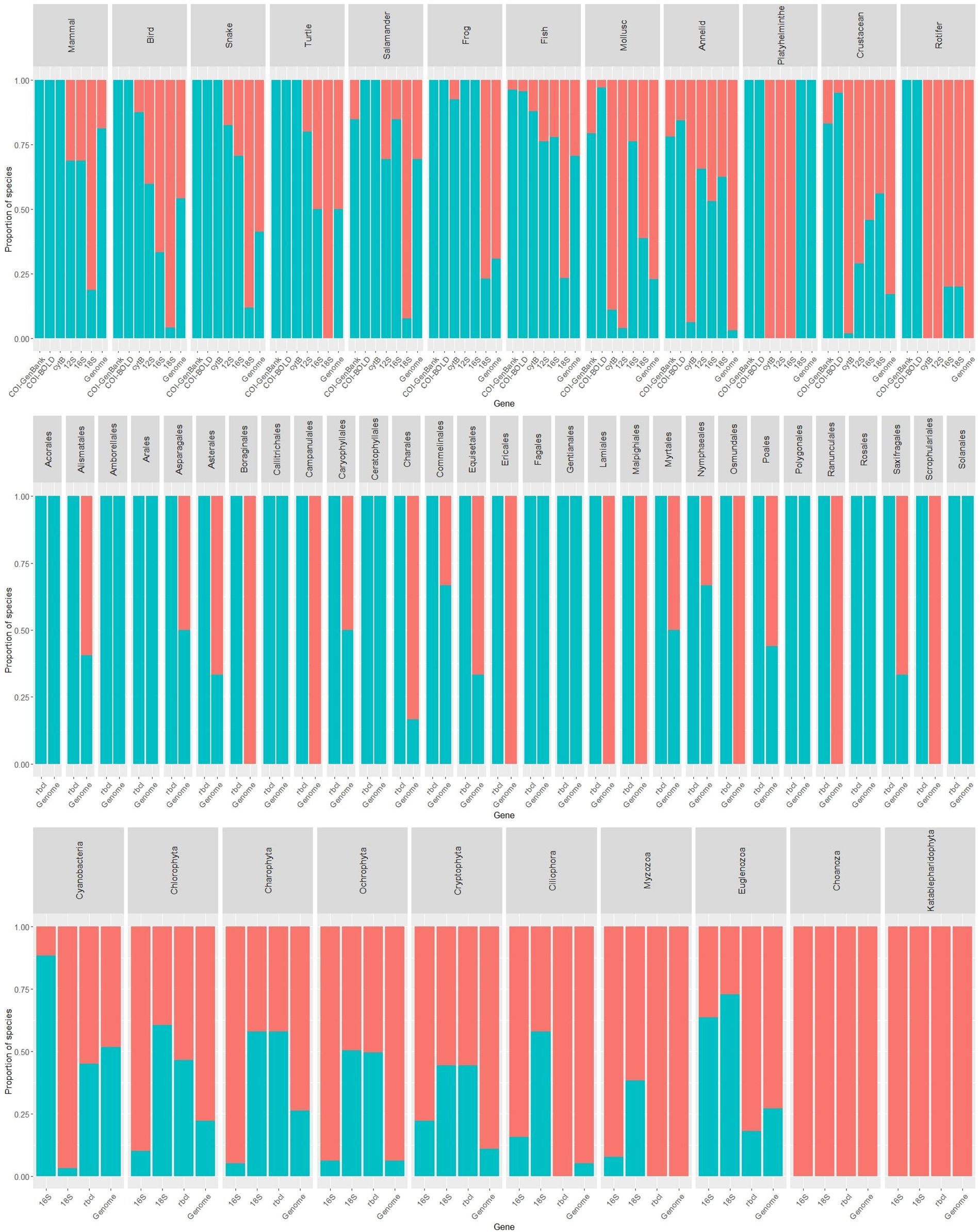
Sequences availability of the species from Eastern Ontario-Western Quebec in the Genbank (COI, cytB, 12S, 16S, 18S, rbcl, genome) and BOLD (COI) public reference databases for A) animals, B) plants, and C) phytoplankton groups. Blue indicates the presence of sequences and red indicates the absence of sequences in the reference database for the corresponding gene. Species used to make this plot are detailed in Table S6.

#### 1.2. Evaluation in silico of the primer pairs

Overall MG2 (COI) and B2 (COI) had the lowest mean penalty scores for vertebrates (mammals, birds, reptiles, amphibians, fishes) and insects (MPS_MG2_ and MPS_B2_ < 100) and are therefore expected to perform better than the other primers pairs for these groups of taxa (Figure 2). Surprisingly, the insect-specific fwhEPTD primer pair showed higher penalty scores (73.0 < MPS < 395.1) than primer pairs not specific to insects, especially for Trichoptera (MPS = 395.1), Ephemeroptera (MPS = 187.2) and Odonata (MPS = 151.9) (Figure 2). MG2 (COI), Leray (COI) and B2 (COI) had the lowest mean penalty scores for crustaceans and rotifers (Figure 2). 16SMOL and 16SMOL2 had the lowest penalty scores for the mollusc and platyhelminth orders (Figure 2). Uniorbcl always had lower MPS for plants compared to its original version, orbcl2, except for Caryophyllales (MPS _Uniorbcl_ = 134.5; MPS _orbcl2_ = 18.7) (Figure 2). Uni16S (16S) was expected to perform very well for cyanobacteria (MPS < 20) and both 18S primer sets were expected to performed well on most eukaryotic phytoplankton orders, although overall UniPh (18S; 0 < MPS <91) was expected to perform best than its original version Uni18S (0 < MPS < 274.6) (Figure 2).

**Figure 2.**
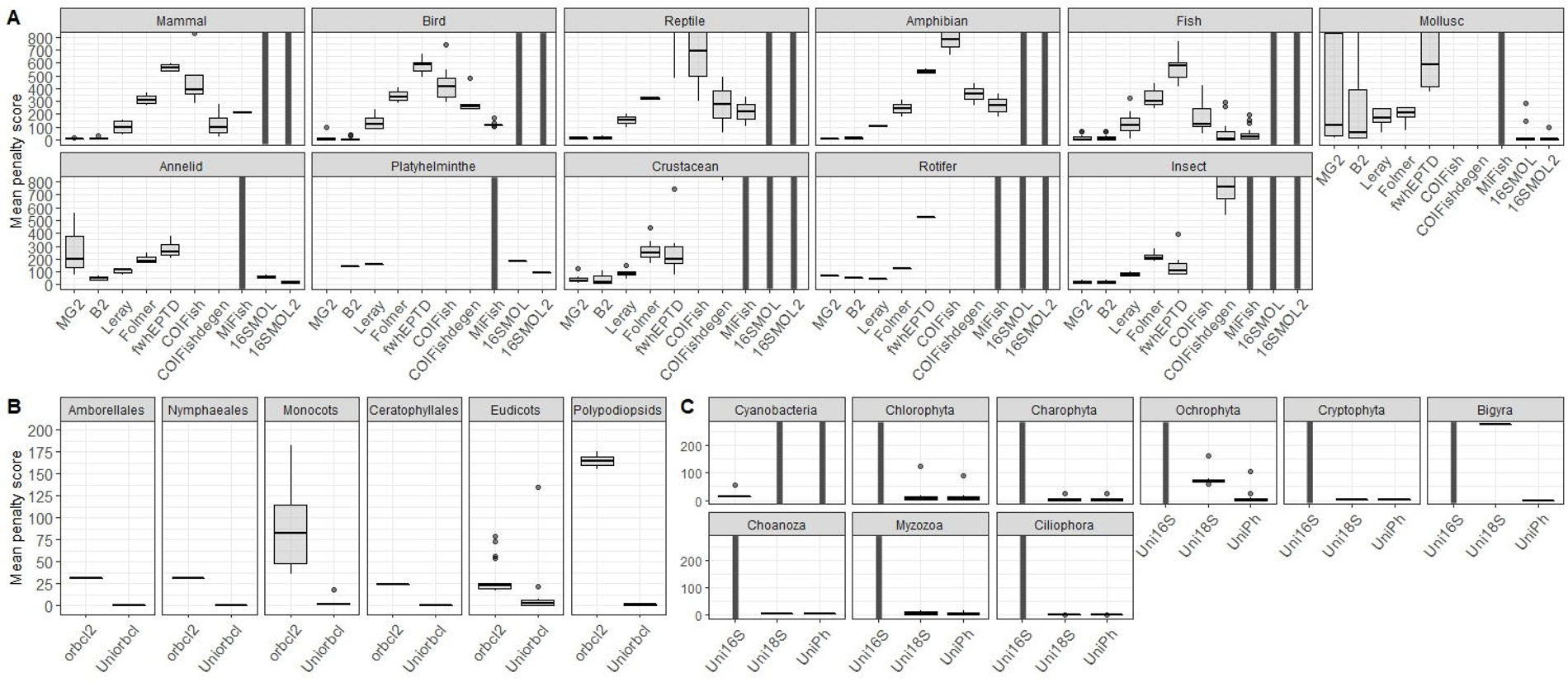
Penalty scores for each primer set, with dots indicating the mean values of each order for A) animals, B) plants, C) phytoplankton. The higher the penalty score calculated using PrimerMiner, the worse the primer is expected to perform. Taxa showing penalty scores higher than 800 for animals (min = 0, max = 5604.1, median = 152.5) and 200 for phytoplankton (min = 0, max = 274.6, median = 3.5) are not represented, on the figure A and B respectively, for better visualisation of lower scores. A vertical black bar indicates that this group was not analysed for the corresponding gene.

### 2. *In vitro* and *in vivo* evaluation: mock communities and eDNA samples

#### Sequencing and filtering results

Our Illumina sequencing produced 17,945,331 reads total of which 16,715,207 passed Illumina filters with 12,014,198 assigned to the samples using the index combination (Table S7). We retained 2,096,942 reads after processing and annotating the sequences in barque and a total of 1,750,554 reads after filtering the output data from barque: 1,538,507 reads in the mock communities (expected species: 1,100,086 reads; details by primer in Table S7), 58,391 reads for the aquarium tanks (expected species: 14,640 reads; details by primer in Table S7), and 136,618 for the eDNA samples (details by primer in Table S7). Note that the Folmer primer pair was not processed because the gap between R1 and R2 was too large.

#### 2.1. Mock communities

Thirty-six taxa (six phytoplankton, 17 plants, two fish, one snake, one insect, four zooplankton and four molluscs) out of 145 were not recovered by any of the primer sets in MC_1_ (11/91), MC_2_ (4/73), MC_3_ (1/8), MC_4_ (6/9), and MC_5_ (29/61) (Table S3). Some taxa with challenging morphological identification were successfully identified at the species level based on their barcode (e.g., *Moxostoma anisurum*, *Ceratophyllum echinatum, Gerris insperatus*, *Sphaerium striatinum*). Conversely, some taxa identified at the species level based on morphology were recovered only at the genus/family level based on their barcode (e.g., *Elodea* sp., *Potamogeton* sp., *Typha* sp.). Taxonomic identifications were mostly consistent across (Table S3) and within genes except for *Pseudacris maculata* in MC_5_ (identified as *P. triseriata* by all COI primer pairs but MG2), *Elliptio complanata* in MC_1_ and MC_3_ (identified as *E. crassidens* by MG2) and *Branta canadensis hutchinsii* in MC_1_ and MC_2_ (identified as *B. hutchinsii* by Leray) (Table S3). The overall percentage of expected taxa identified at the species level was influenced by the gene (F_df=5_ = 23.67, *p* << 0.001) but not the length of the target fragment (*p* = 0.34) nor the percentage of degenerate bases (*p* = 0.99). The percentage of taxonomic identification at the species level was higher for COI than for r16S (*p* = 0.03), 18S (*p* = 0.02) and rbcl (*p* < 0.0001), and higher for m16S than for r16S (*p* = 0.02), 18S (*p* = 0.01), and rbcl (*p* = 0.003).

The COI B2, Leray, and fwhEPTD primer pairs had ≥10 detections discarded because of replicate- and negative control-based filtering (Table S3). Overall MG2 recovered the highest number of taxa (N=99) and had the highest number of detections (161 detections at the species level, four at the genus level and two at the family level), followed by Leray (89 taxa, 151 detections at the species level and one at the genus level) and B2 (88 taxa, 148 detections at the species level and two at the genus level) (Figure 3 and Table S3). In MC1, Leray recovered the highest number of invertebrates (7/8), MG2 the highest number of vertebrates (64/73), Uni16S the highest number of phytoplankton (1/1) and orbcl the highest number of plants (3/8). In MC2, MG2 recovered the highest number of taxa (64/73) while Leray and MG2 were equivalent to MC3 (7/8). In MC_4_ the best primer pairs were 16SMOL, 16SMOL2 and Uni16S for phytoplankton (1/1), and Uniorbcl and orbcl for plants (3/8). Finally, MG2 was the best in MC_5_ for invertebrates and vertebrates (9/18 and 18/19 respectively) while UniPh, orbcl and Uni16S were the best for phytoplankton (1/6) and Uniorbcl and orbcl for plants (7/18). We observed a significant effect of the number of reads (F_df=1_ = 83.55, *p* << 0.001) and the primer degeneracy (F_df=1_ = 4.88, *p* = 0.030) on the proportion of detected taxa. We run the same analyses without the primer pairs counting less than 5% of the total reads (< 55,005 reads) to account for potential outliers driving the signal without losing too much information (McMurdie and Holmes 2014): we observed an effect of the number of reads (F_df=1_ = 47.243, *p* << 0.001) and the length of the target fragment (F_df=1_ = 4.571, *p* = 0.04), but no effect of primer degeneracy (*p* > 0.05).

**Figure 3.**
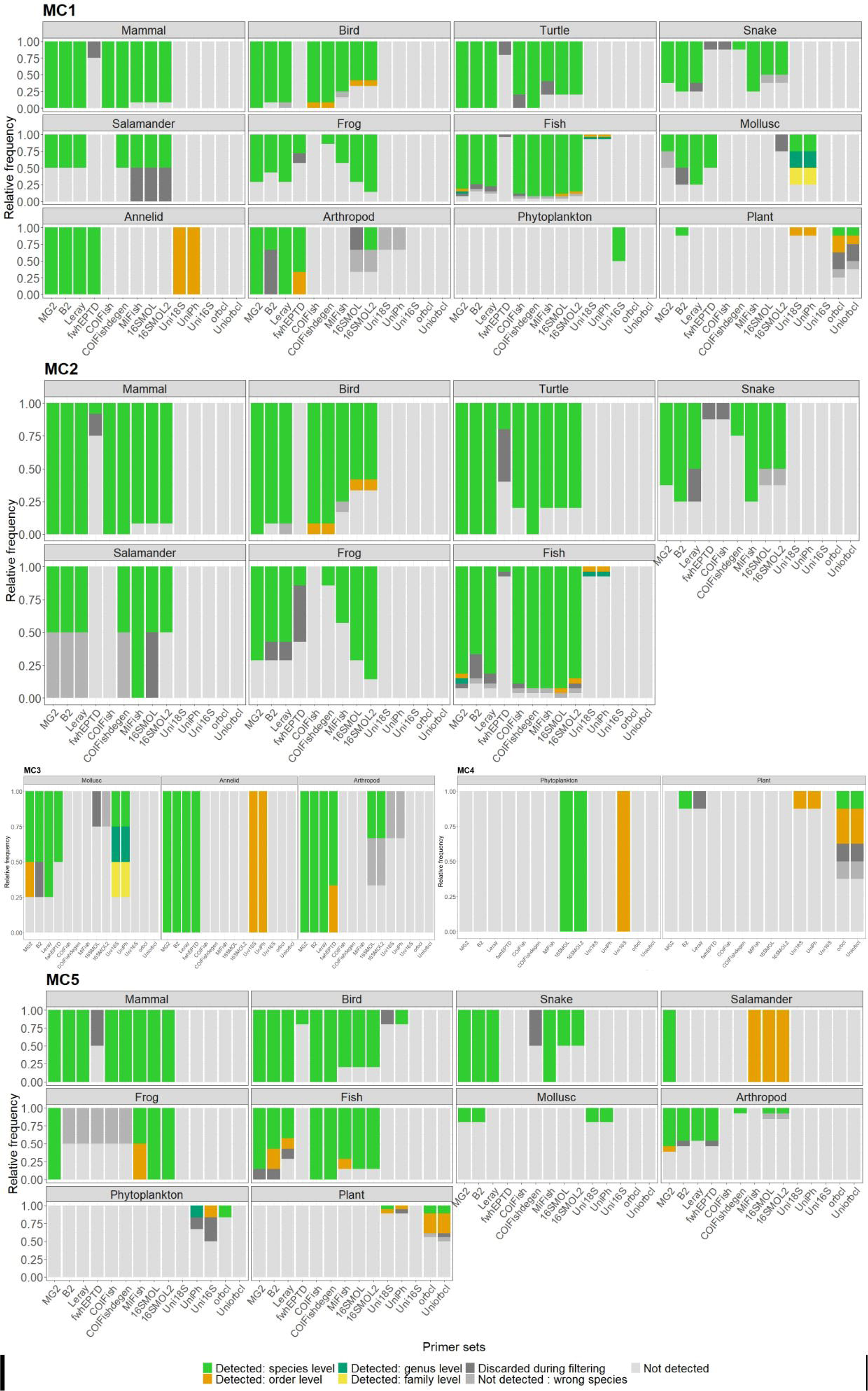
Relative frequency of the detections for each primer pair in the five mock communities. “Discarded during filtering” groups both inconsistency among replicates and negative control-based filtering. “Not detected: wrong species” indicate that a taxon as detected but the species identification was likely wrong (e.g., *Ranatra australis* instead of *Ranatra fusca*).

Taxa included in more than one mock community showed similar detection success among them (i.e., same primer pairs detecting the taxa at the species level in the mock communities). For example, the fish *Ambloplites rupestris* was detected by the same eight primer pairs in both MC1 and MC2. Overall, 20 taxa (one mollusc, two arthropods, one phytoplankton and 16 vertebrates) showed differences in detections among mock communities (i.e., difference in the number of primer pairs detecting the taxa, Figure 4) because of inconsistent technical replicates that were discarded in the filtering process (Table S3).

**Figure 4.**
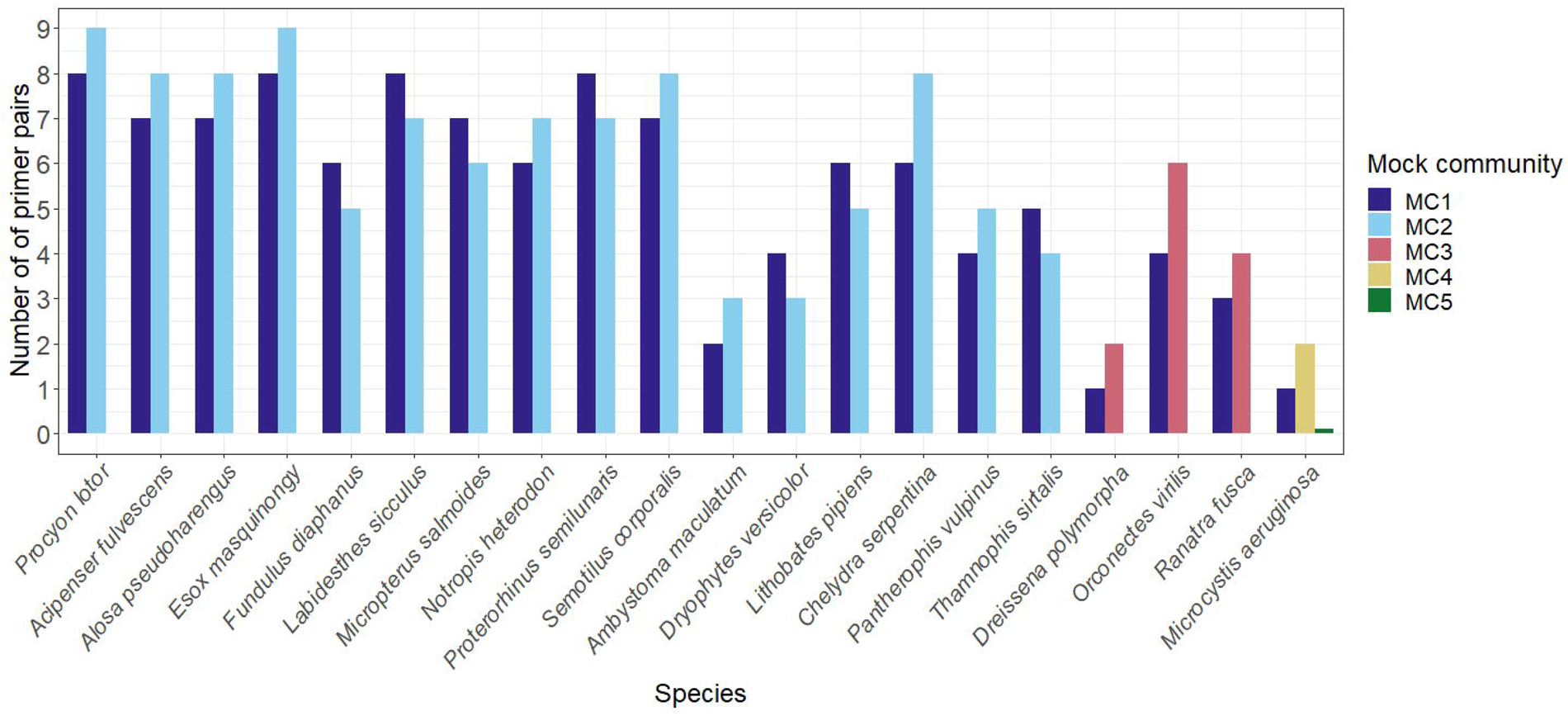
Inconsistencies (number of primer pairs) of species detection between mock communities.

#### 2.2. Aquarium tanks

The eDNA extraction for the first aquarium tank (BA1) failed so none of the species were recovered in this sample. However, the red belly dace (*Chrosomus eos*) from BA1 was surprisingly identified in the second tank (BA2). We recovered some taxa that were part of the fish diet (e.g. *Mysis*, shrimp, rainbow smelt, poultry and other protein meals), turtle/frogs diet (e.g. cricket, worms) and otter diet (e.g. mackerel, hake) (Table S4), as well as microscopic taxa such as water mold, or acari that are common of the natural materials in the terrariums.

We identified 62.5% of the 24 species present in the six other tanks, all to species level except *Lepisosteus osseus* identified to genus by MiFish (Table S4). We did not observe any effects of gene, target amplicon length or degenerate bases on the percentage of taxa identified at the species level (*p* > 0.3). Five species (20.8%) were detected but did not pass our conservative filtering steps (two positive replicates and/or threshold of three reads minimum): the rockbass (*Ambloplites rupestris*), the channel catfish (*Ictalurus punctatus*), pumpkin seed (*Lepomis gibbosus*), blue gill (*Lepomis macrochirus*) and spotted turtle (*Clemmys guttata*) (Table S4). Two fish species from BA2 were misidentified by our markers: the striped bass (*Morone saxatilis*) was identified as the white bass (*M. chrysops*) and the fallfish (*Semotilus corporalis*) was identified as the creek chub, *S. atromaculatus* (Table S4). Two species were not detected by any primer pair despite being identified in our eDNA samples taken from natural systems: the spot fin shiner (*Cyprinella spiloptera*), and yellow perch (*Pomoxis nigromaculatus*). Neither the number of reads (*p* = 0.06), nor the other explanatory variables (*p* > 0.25) had significant effects on the proportion of detected taxa.

#### 2.3. eDNA samples from natural systems

We obtained a total of 461 taxa from 34 phyla, 135 orders, 193 families and 294 genera over the six eDNA samples (Figure 5), including 22 taxa either: endangered, invasive, introduced, parasite, pest, toxic, or water quality indicator (Table 2).

**Figure 5.**
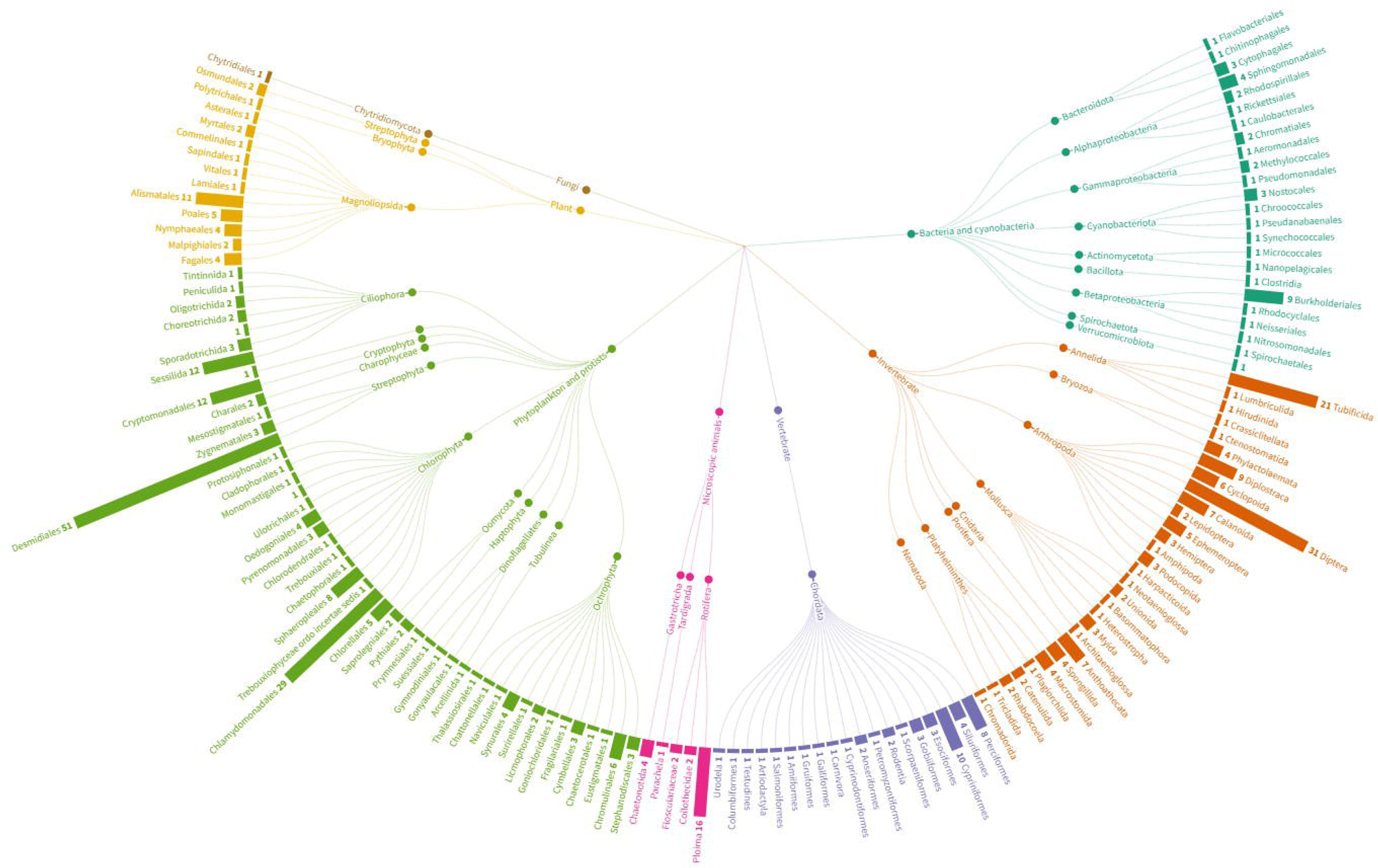
Taxonomic phylogram of the range of taxa detected in the natural systems. The figure was made using Flourish Studio App (https://app.flourish.studio/).

Overall, the rbcl and 18S primer pairs identified a higher number of taxa but with poorer resolution: an average of 64.6% and 72.8% of taxa identified at the species level respectively, versus 85%, 100% and 92% for the COI, 12S and mitochondrial 16S primer pairs respectively (Figure 6). Note that we manually adjusted the bacteria identification at the genus level, so the Uni16S resolution percentage was underestimated, and it was therefore not included in the resolution analyses. The percentage of taxa identified at the species level was influenced by the gene (F_df=4_ = 7.797, *p* << 0.001), with a higher percentage for COI than 18S (*p* = 0.007) and rbcl (*p* < 0.002). We observed only a marginal effect of the length of the target fragment (*p* > 0.053) and no effect of the percentage of degenerate bases (*p* > 0.707).

**Figure 6.**
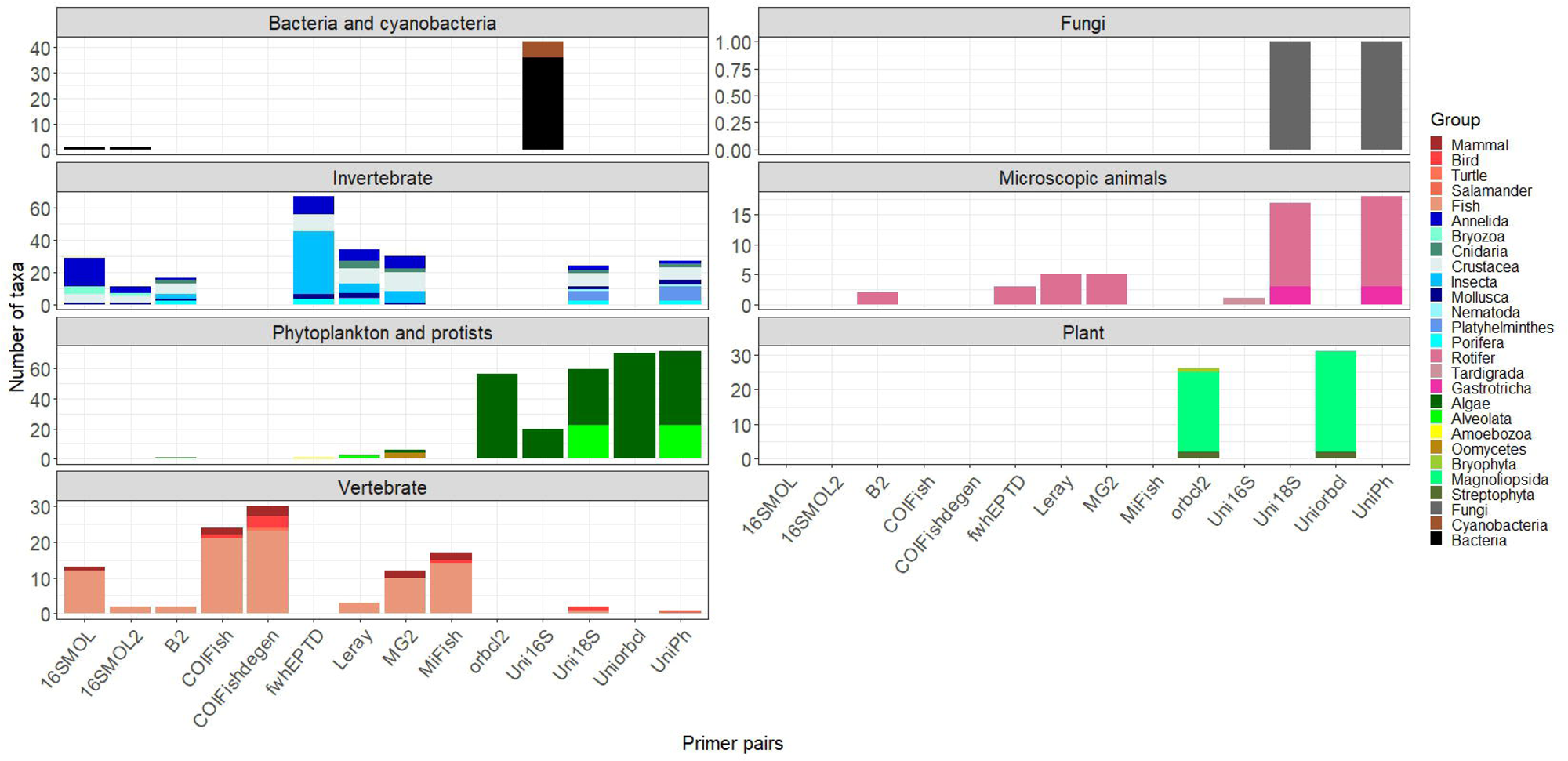
Number of taxa detected by each primer pair in the natural systems.

The highest number of vertebrates was recovered using COIFishdegen (N=30), invertebrates using fwhEPTD (N=67), phytoplankton and protists using UniPh (N=71), plants using Uniorbcl (N=31) and bacteria using Uni16S (N=36) (Figure 6). The number of taxa was positively impacted by the number of reads (F_df=1_ = 63.594, *p* << 0.001), the length of the target fragment length (F_df=1_ = 8.150, *p* = 0.005), and the percentage of degenerate bases (F_df=1_ = 8.114, *p* = 0.005). We found the same results when running the same analyses without the primer pairs counting for less than 1.5% of the total reads (< 2000 reads; 16SMOL2 = 440 reads, MiFish = 1714 reads, B2 = 1976 reads).

#### 2.4. Decision tool for selecting primer pairs

We created a free online tool – SNIPe – that allows users to explore eDNA data for multiple primer pairs and select combinations that best serve the goals of their study (https://snipe.dlougheed.com). Applied to our eDNA dataset, analyses using SNIPe indicated that 13 primer pairs would be required to recover 100% of the taxa. Using our best single primer pair (UniPh), we recovered 25.6% of all taxa. Thereafter, combining our two best primer pairs (UniPh + Uniorbcl), we recovered 46.9% of all taxa; for three (UniPh + Uniorbcl + fwhEPTD), 61.8%; for four, 74.8% (UniPh + Uniorbcl + fwhEPTD + Uni16S); and for five (UniPh + Uniorbcl + fwhEPTD + Uni16S+16SMOL), 82% We illustrate the user interface and workflow for SNIPe in Figure 7.

**Figure 7.**
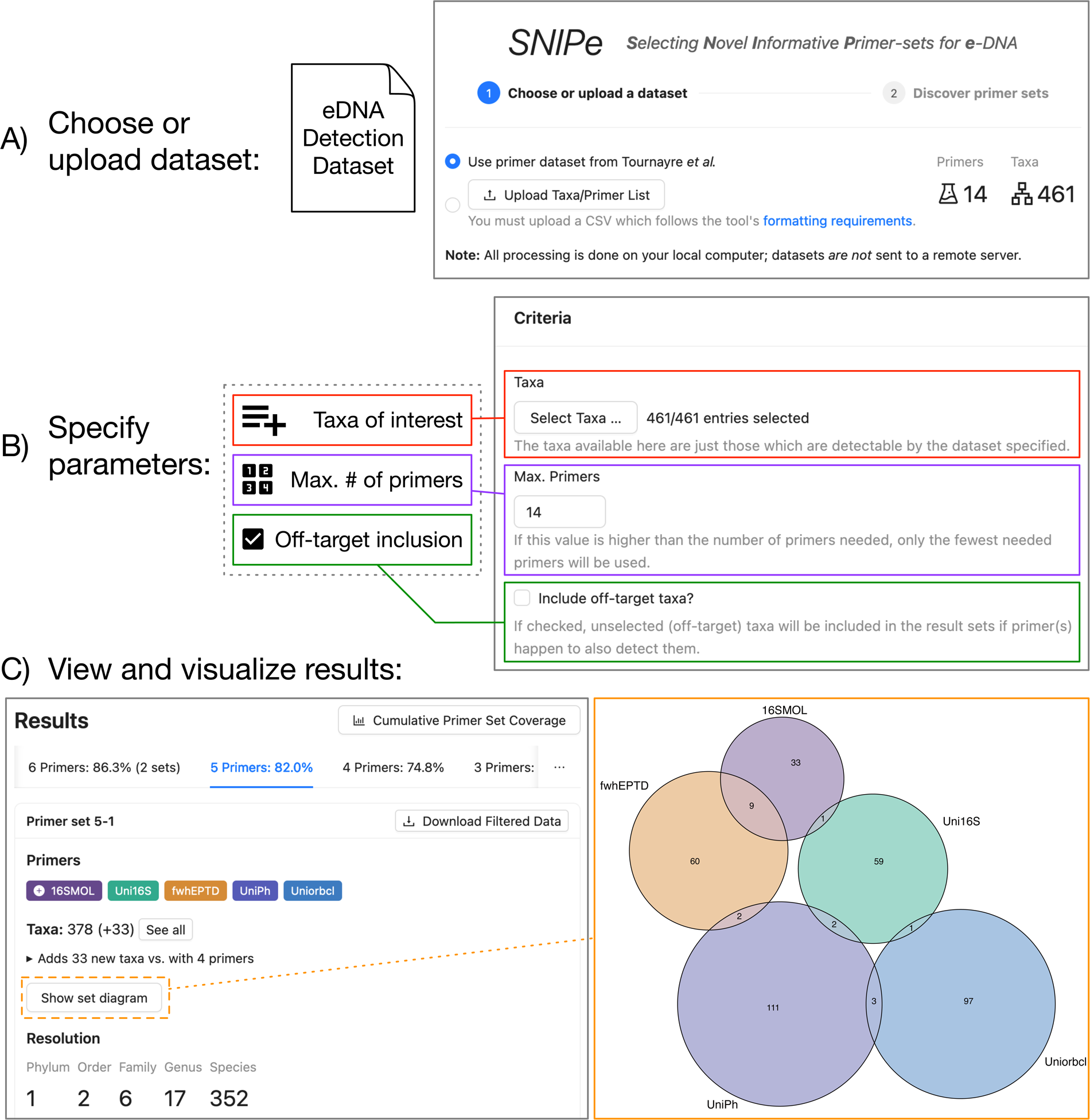
Workflow and user interface for the SNIPe tool. Given A) a presence-detection dataset, and B) a set of taxa of interest and any additional desired parameters, a range of primer sets with different numbers of primer pairs are computed. The best primer set(s) for each number of primers are reported to the user (C). Users can then examine the primer set resolution and taxon coverage and visualize primer pair coverage overlap.

## DISCUSSION

### 1. Describing freshwater communities

Our study highlights the potential for eDNA-based methods to quantify aquatic biodiversity, monitor rare and cryptic species, invasive species, and assess trophic interactions (Beng and Corlett 2020; Bohmann et al. 2014). We were able to detect hundreds of species across the tree of life, including several species of interest relevant to conservation, human health and agriculture: model species in toxicity testing (e.g. Diptera, Annelida), species at risk (e.g. the Paper pondshell, *Utterbackia imbecillis*, a species at risk mollusc in Canada), pathogens (oomycete, toxin producing algae), parasites (e.g. parasitic fish, chytrid), pests (e.g. lepidoptera Spongy moth, diptera Spotted-wing *Drosophila*), and invasive species (e.g., fish: Round and Western tubenose gobies, Sea lamprey; molluscs: Zebra and Quagga mussels; plant: Purple loosestrife). The use of eDNA as a method for the cost-effective and non-intrusive monitoring and management of invasive species and species at risk has seen increasing attention and standardization in recent years (Abbott et al. 2021; Fediajevaite et al. 2021). The concomitant detection of invasive/native species can help to delineate the invasion front of an invasive species (Keller et al. 2022) and capture shifting dynamics of the river ecosystem (Greenhalgh et al. 2022), for example the increasing detection in time and space of dreissenid mussels to the detriment of local native species (Ricciardi, Whoriskey, and Rasmussen 1996).

Our study also showcases the power of water eDNA-based metabarcoding to reconstruct freshwater communities in connected systems. For example, we were able to detect species from different tanks by collecting water samples from the shared water system and detect taxa that comprised the diet of some captive species. While some unexpected detections could be attributed to wrong species identification by the primers (e.g. stripe bass, fall fish), the detection of the red belly dace in the Fresh Water Fish water system (BA2) was surprising because it has been a long time since they were in that system, and it was only for a short period of time. As DNA detectability in the water column is limited to a few weeks (Dejean et al. 2011), and water at this facility is treated, the source of this unexpected detection is likely contamination during sampling.

Although we cannot rule out the possibility of contamination from the mock communities or DNA in the lab despite conservative filtering, our results would be consistent with eDNA transport by water flow from upstream lakes into the marshes and ponds (Deiner and Altermatt 2014; Laporte et al. 2020; Wacker et al. 2019). For example, we detected the yellow perch (*Perca flavescens*) within a shallow palustrine marsh that cannot be inhabited by this species, but the marsh does connect to a small freshwater lake upstream where the presence of the species is known. While such aquatic transport of eDNA may be considered an issue for deducing location of a detected species or local species richness, it can also be used to detect cryptic species and make powerful inferences in species distributions. For example, Deiner et al. (2016) showed that eDNA data from sampling stream networks can provide insights on biodiversity distribution across hierarchical spatial scales, including both terrestrial and aquatic species. Many species detected with eDNA metabarcoding were known to occur at the sampling sites based on land-based sampling (BIObus Canadian National Parks Malaise Program), or traditional aquatic surveys (Queen’s University Biological Station, Thousand Islands National Park and the River Institute). Interestingly, some expected and rather abundant species, such as the American leech (*Macrobedella decora*) or Broadleaf arrowhead (*Sagittaria latifolia*) were not recovered in some of the eDNA samples. The false negative detections could be due to several factors: an insufficient sampling coverage (Guillera-Arroita et al. 2017), competition by DNA templates of other taxa during PCR (Kelly, Shelton, and Gallego 2019; Marinchel et al. 2023), or our conservative filtering approach (Alberdi et al. 2018; Mathon et al. 2021). Thus, we recommend having taxonomists and ecologists check the lists of species obtained from eDNA data as this can reveal both potential false positives and false negatives, and guide interpretation of the results. A synergistic approach combining traditional surveying techniques and Indigenous knowledge with eDNA can validate otherwise weakly substantiated eDNA datasets (Lopez et al. 2023).

### 2. Validation and selection of the best primer pair combination(s)

Our study highlighted disparities in the completeness of public databases of reference sequences across the tree of life and among genes. Despite duplication between databases (Meiklejohn, Damaso, and Robertson 2019), our analysis also showed differences in species coverage between databases for the same gene: the BOLD database encompassed more species than GenBank for COI, potentially due to the dramatic expansion in geographic and taxonomic coverage in BOLD by the international Barcode of Life consortium (BIOSCAN program, Hobern 2021). While we were able to recover most animal and plant species, we observed a lack of reference sequences for phytoplankton in GenBank (∼40% of the species included in our study did not have sequences for any of the tested genes) probably because of the challenge of phytoplankton morphological identification to the species level (McManus and Katz 2009; Tzafesta et al. 2022). Similar to other studies, we observed differences between genes (e.g., COI covered more mammal species than 18S in GenBank) (Marques et al. 2021) and metazoan species without COI reference sequences had no sequences available for any other marker. This favors the use of COI primers for animal metabarcoding studies, yet it is not the ‘gold standard’ for all animal taxonomic groups. For example, most fish metabarcoding studies are standardized with primer pairs for a portion of the mitochondrial 12S gene (e.g., MiFish, Masaki Miya, Gotoh, et Sado 2020; teleo, Valentini et al. 2016), partly because 12S primer binding regions are less variable than those for COI and thus they co-amplify less non-fish species (Macher et al. 2023; Stat et al. 2017). There is then a key need for completing 12S general and fish specific databases such as MitoFish (Iwasaki et al. 2013; Sato et al. 2018; Collins et al. 2019; Macher et al. 2023).

Reference sequences are critical for evaluating primer pairs on a large scale as we have attempted: they are used *in silico* to predict amplification success by calculating a score based on mismatches between primer and template sequences (Ficetola et al. 2010; Elbrecht et Leese 2016). Such analysis can include consideration of the number of mismatches, their positions on the sequence and the adjacency of mismatches. However, *in silico* results do not always match *in vitro* results (e.g. Tournayre et al. 2020). First, the sequences contained within reference databases are produced using different approaches (e.g., Sanger Sequencing, whole genome shotgun sequencing) and different primers pairs may target different parts of a focal gene. This can result in heterogeneity in the availability and quality of the sequences, and in species recovery among primer pairs for the same gene. Second, PCR conditions can greatly influence the performance of primers (e.g., annealing temperature, number of cycles) (Krehenwinkel et al. 2017) and *in silico* analyses cannot predict PCR bias (e.g., DNA competition, amplification stochasticity). Intraspecific variation, particularly regional variation, can also limit *in silico* primer testing (Estensmo et al. 2021). In our study, *in silico* and mock community results were very similar for metazoans: overall the COI mini-barcode MG2 (Tournayre et al. 2020) was the best for invertebrates and vertebrates, closely followed by Leray (Leray et al. 2013) and B2 (Elbrecht and Leese 2017). Contrary to the expectations from the *in silico* analyses, the 16S mollusc primers (16SMOL, Klymus, Marshall, et Stepien 2017; and 16SMOL2, this study) did not recover more molluscs than the other primer pairs, potentially because we did not use blocking primers that would have favored mollusc amplification (Klymus, Marshall, et al. 2017). It is especially likely for 16SMOL2 because these primers have a high proportion of degenerate bases within primers (∼26%) that probably increase the amplification of non-target species. Surprisingly, Uniorbcl, our degenerate version of the orbcl2 plant primer (Coghlan et al. 2021), and UniPh, our degenerate version of the Uni18S primer pair (Zhan et al. 2013) did not perform better than their original version in mock communities. However, they did appear to perform best in the eDNA samples from our sampled water bodies, so the limited taxonomic range of the mock communities may have limited the potential of these primer pairs. Such disparities in interpretation emphasize the usefulness of combining controlled experiments (*in silico*, mock communities, aquarium) and field surveys to assess the benefits and the limitations of primer pairs before conducting full scale metabarcoding studies. Similarly to the species-specific assay validation scale proposed by Thalinger et al. (2021), the metabarcoding primers evaluation would benefit from a similar validation schema (Figure 8).

**Figure 8.**
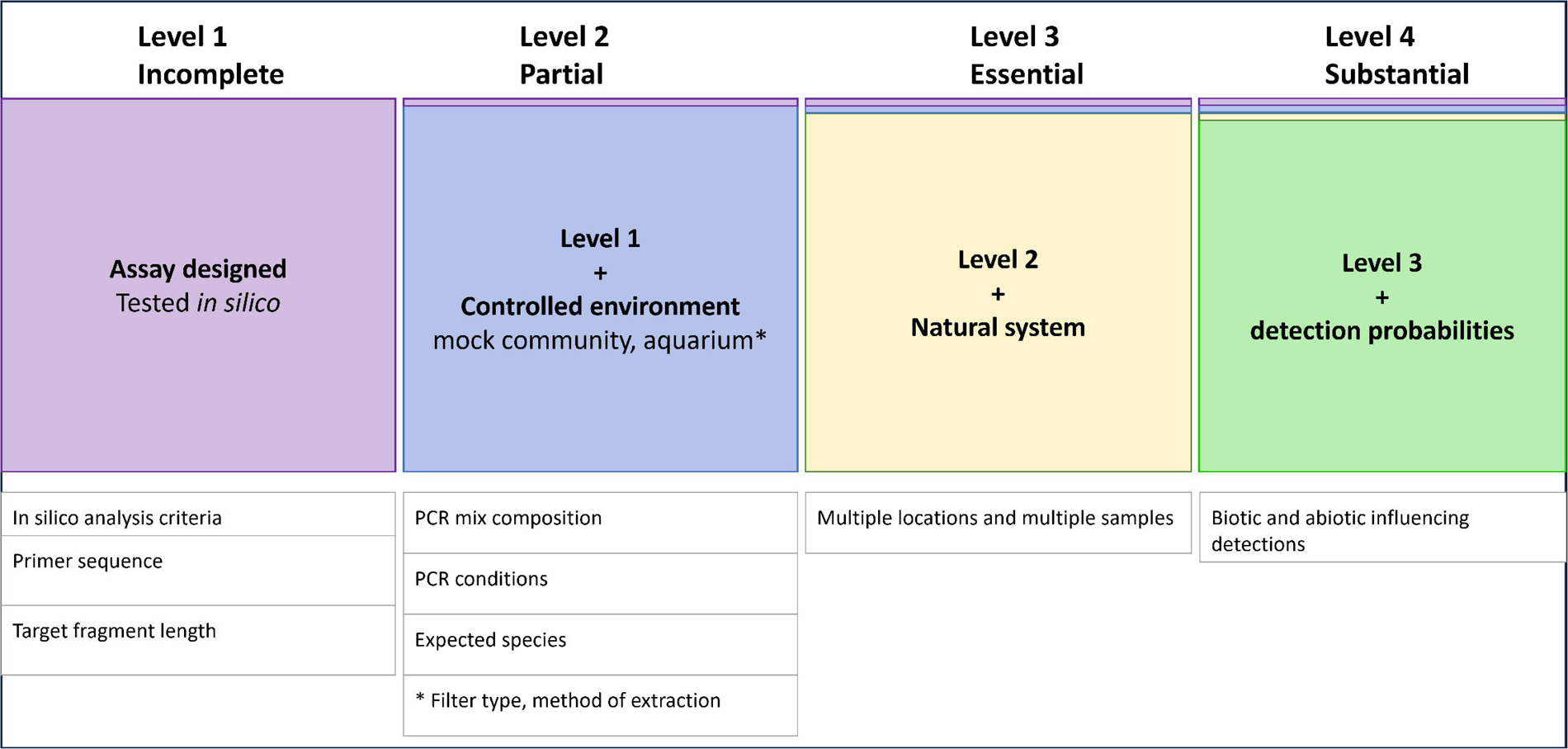
Primer validation scale and minimum required information for metabarcoding studies.

### 3. Maximizing prospects for successful metabarcoding

Filtering steps and their stringency can greatly influence detection success. For example, we showed that our conservative filtering approach (“true detection” = two positive PCR replicates out, three or more reads) decreased the risk of false positives at the cost of increasing the probability of false negatives. PCR replicates are important to account for PCR and sequencing stochasticity, and primer efficiency, especially in eDNA samples which have low and heterogeneous DNA concentrations (Ficetola et al. 2015). However, because rare taxa are recovered stochastically they are often unique to one PCR replicate and would be considered as a false positive (Shirazi, Meyer, and Shapiro 2021). Increasing sequencing depth to recover more taxa also concomitantly increases the dissimilarity among PCR replicates by increasing the number of sequences originating from contamination and chimeras (Alberdi et al. 2018; Deagle et al. 2014). However, sequencing depth is important, although our consideration of this variable was limited in our study because we used a single sequencing run for all sample types and primer pairs. Pooling replicates before indexing could have been done but this is not necessarily more efficient for assessing diversity and community structure (Smith et Peay 2014); for example, while pooling reduces the risk of missing taxa it also artificially inflates the possibility of false positives due to accumulation of artefactual sequences (Alberdi et al. 2018). Thus, we emphasize the need for clarity and transparency in the methods and results of metabarcoding studies for relevant interpretations of the data.

Primer amplification success was influenced only by primer degeneracy in the mock communities (where DNA was not degraded), and by both primer degeneracy and the length of the target fragment in the eDNA samples. Primer degeneracy minimizes mismatches between primers and template sequences for a range of taxa, but the length of the target fragment is also crucial because long fragments are rarer than short ones in water (Bylemans et al. 2018) and eDNA will likely break into smaller fragments over time. In sum, our results support using degenerate primers and short amplicons if the goal is to amplify as many species as possible while minimizing the number of eDNA metabarcoding primer pairs. Independent of considerations of target length and primer degeneracy, the choice of gene is pivotal as it influences the taxonomic resolution (i.e., the taxonomic level at which taxa can be identified): as in other studies (Deagle et al. 2014; Stat et al. 2017), our results show that nuclear markers provided a broad taxonomic coverage (e.g. 18S rRNA) but less resolution than mitochondrial ones (e.g., COI, 12S). A lack of reference sequences can lead to wrong identification with seemingly high confidence. For example, 16S rRNA primers identified the wrong *Ranatra* species with high confidence (>98% identity) because *Ranatra fusca* is absent from the database. Moreover, a lack of reference sequences can have a proportionately greater impact on taxonomic resolution for short compared with long amplicons. The chance of getting a species identification with high confidence (high %identity and %coverage) for the wrong taxa is higher (Tournayre et al. 2020). Incorrect molecular identification can also result from complex phylogenetic relationships or changing systematics and taxonomic synonyms that lead to errors in reference databases. For example, the taxonomy of the western chorus frog (*Pseudacris triseriata*) is controversial in our study area because of apparent mito-nuclear discordance and the presence of two deeply divergent mitotypes (*Pseudacris triseriata/maculata*) (Lougheed et al. 2020). Finally, it is also possible that some plants in our mock communities were not detected because of morphological misidentification as they were not checked by plant experts. For example, the species we had identified based on morphology as *Juncus effusus* (MC4 and MC5), was detected with a strong signal as *Eleocharis* sp. instead (Uni18S: 8839 reads, UniPh: 10228 reads, orbcl: 8313 reads and Uniorbcl: 6508 reads) which is phenotypically similar.

A challenge when designing an eDNA study is selecting combinations of primer pairs which best serve the objectives of the study. We developed our online tool, SNIPe, to help researchers focus on a particular set of taxa relevant to their study (e.g., predator and prey, competing native and invasive species, or species at different trophic levels), and to optimize which, and how many, primer pairs they should use to obtain occurrence data to best address their research questions. We provide a default dataset comprising 14 primer pairs and the suites of species detected for each. SNIPe can also accommodate user-supplied datasets from work in their own labs, or data compiled from literature, allowing for powerful cross-study primer selection. All dataset processing and computation is done on the users’ own computer inside the web browser, meaning no user-uploaded data is transmitted to nor retained on the SNIPe web server.

## CONCLUSION

In conclusion, our workflow combining in silico, mock communities, aquarium and eDNA samples from natural systems, showed that multi-marker DNA metabarcoding is very promising for the evaluations of entire freshwater biotic communities. Our results revealed that even with a low number of combined primer pairs, it is possible to identify taxa from primary producers through herbivores to predators, native and invasive species, and rare and common species. As the choice of the primers (target taxa, gene, length of the amplicon, degeneracy) is pivotal for the success of metabarcoding studies, we provide an online tool that will guide primer selection based on the goal of the study and the trade off species coverage-cost. Ultimately, our study represents a step forward in overcoming the technical challenges associated with the holistic monitoring of freshwater communities.

## AUTHORS CONTRIBUTION

Design of the study: OT, SCL

Field work: OT, HT, MJSW

Lab work: OT, HT, ZS

Bioinformatics: OT, DRL

Analysis: OT

Interpretation of the data: OT, SCL, MJSW, SL, JC

Drafting of the manuscript: OT, SCL

Editing of the manuscript: all authors

Funding acquisition: SCL, BFC, SEA, YW, JR

## DATA ARCHIVING STATEMENT

The MiSeq raw data (R1 and R2 .fastq.gz files) and indexing sheet were archived in the Figshare public repository: https://figshare.com/s/71fa7c10cb0e0ac94a6a

## ACKNOWLEDGMENT

We thank Burton Lim, Maureen Zubowski, Amy Lathrop and Santiago Claramunt (Royal Ontario Museum), and Barb Vanderbeld (Queen’s University) for providing samples for the mock communities. We are grateful to Jenna Shannon and Lena Campos for helping with DNA extraction. This work was funded by a SSHRC New Frontiers in Research Fund – Exploration (NFRFE) grant, an NSERC Alliance grant, a South Frontenac grant, and the Baillie Family Chair in Conservation Biology.

## REFERENCE

Abbott, Cathryn, Mark Coulson, Nellie Gagné, Anaïs Lacoursière, Robert Bajno, Charise Dietrich, and Shannan May-McNally. 2021. “Guidance on the Use of Targeted Environmental DNA (eDNA) Analysis for the Management of Aquatic Invasive Species and Species at Risk.” DFO Can. Sci. Advis. Sec. Res. Doc. 2021/019. *Iv* +42p.

Abell, Robin, Michele L. Thieme, Carmen Revenga, Mark Bryer, Maurice Kottelat, Nina Bogutskaya, Brian Coad, Nick Mandrak, Salvador Contreras Balderas, William Bussing, Melanie L. J. Stiassny, Paul Skelton, Gerald R. Allen, Peter Unmack, Alexander Naseka, Rebecca Ng, Nikolai Sindorf, James Robertson, Eric Armijo, Jonathan V. Higgins, Thomas J. Heibel, Eric Wikramanayake, David Olson, Hugo L. López, Roberto E. Reis, John G. Lundberg, Mark H. Sabaj Pérez, and Paulo Petry. 2008. “Freshwater Ecoregions of the World: A New Map of Biogeographic Units for Freshwater Biodiversity Conservation.” BioScience 58(5):403–14. doi: 10.1641/B580507.

Ahmed, Shams Forruque, P. Senthil Kumar, Maliha Kabir, Fatema Tuz Zuhara, Aanushka Mehjabin, Nuzaba Tasannum, Anh Tuan Hoang, Zobaidul Kabir, and M. Mofijur. 2022. “Threats, Challenges and Sustainable Conservation Strategies for Freshwater Biodiversity.” Environmental Research 214:113808. doi: 10.1016/j.envres.2022.113808.

Alberdi, Antton, Ostaizka Aizpurua, M. Thomas P. Gilbert, and Kristine Bohmann. 2018. “Scrutinizing Key Steps for Reliable Metabarcoding of Environmental Samples.” Methods in Ecology and Evolution 9(1):134–47. doi: 10.1111/2041-210X.12849.

Albert, James S., Georgia Destouni, Scott M. Duke-Sylvester, Anne E. Magurran, Thierry Oberdorff, Roberto E. Reis, Kirk O. Winemiller, and William J. Ripple. 2021. “Scientists’ Warning to Humanity on the Freshwater Biodiversity Crisis.” Ambio 50(1):85–94. doi: 10.1007/s13280-020-01318-8.

Andres, Kara J., Timothy D. Lambert, David M. Lodge, Jose Andrés, and James R. Jackson. 2023. “Combining Sampling Gear to Optimally Inventory Species Highlights the Efficiency of eDNA Metabarcoding.” Environmental DNA 5(1):146–57. doi: 10.1002/edn3.366.

Angeler, David G., Craig R. Allen, Hannah E. Birgé, Stina Drakare, Brendan G. McKie, and Richard K. Johnson. 2014. “Assessing and Managing Freshwater Ecosystems Vulnerable to Environmental Change.” Ambio 43(1):113–25. doi: 10.1007/s13280-014-0566-z.

Armingohar, Zahra, Jørgen J. Jørgensen, Anne Karin Kristoffersen, Emnet Abesha-Belay, and Ingar Olsen. 2014. “Bacteria and Bacterial DNA in Atherosclerotic Plaque and Aneurysmal Wall Biopsies from Patients with and without Periodontitis.” Journal of Oral Microbiology 6(1):23408. doi: 10.3402/jom.v6.23408.

Beng, Kingsly C., and Richard T. Corlett. 2020. “Applications of Environmental DNA (eDNA) in Ecology and Conservation: Opportunities, Challenges and Prospects.” Biodiversity and Conservation 29(7):2089–2121. doi: 10.1007/s10531-020-01980-0.

Benson, Dennis A., Ilene Karsch-Mizrachi, David J. Lipman, James Ostell, and David L. Wheeler. 2008. “GenBank.” Nucleic Acids Research 36(Database issue):D25–30. doi: 10.1093/nar/gkm929.

Berger, Chloé Suzanne, Cecilia Hernandez, Martin Laporte, Guillaume Côté, Yves Paradis, Dominique W. Kameni T., Eric Normandeau, and Louis Bernatchez. 2020. “Fine-Scale Environmental Heterogeneity Shapes Fluvial Fish Communities as Revealed by eDNA Metabarcoding.” Environmental DNA 2(4):647–66. doi: 10.1002/edn3.129.

Bohmann, Kristine, Alice Evans, M. Thomas P. Gilbert, Gary R. Carvalho, Simon Creer, Michael Knapp Douglas W. Yu, and Mark de Bruyn. 2014. “Environmental DNA for Wildlife Biology and Biodiversity Monitoring.” Trends in Ecology & Evolution 29(6):358–67. doi: 10.1016/j.tree.2014.04.003.

Bolger, Anthony M., Marc Lohse, and Bjoern Usadel. 2014. “Trimmomatic: A Flexible Trimmer for Illumina Sequence Data.” Bioinformatics 30(15):2114–20. doi: 10.1093/bioinformatics/btu170.

Bylemans, Jonas, Elise M. Furlan, Dianne M. Gleeson, Christopher M. Hardy, and Richard P. Duncan. 2018. “Does Size Matter? An Experimental Evaluation of the Relative Abundance and Decay Rates of Aquatic Environmental DNA.” Environmental Science & Technology 52(11):6408–16. doi: 10.1021/acs.est.8b01071.

Coghlan, Stephanie A., Aaron B. A. Shafer, and Joanna R. Freeland. 2021. “Development of an Environmental DNA Metabarcoding Assay for Aquatic Vascular Plant Communities.” Environmental DNA 3(2):372–87. doi: 10.1002/edn3.120.

Collins, Rupert A., Judith Bakker, Owen S. Wangensteen, Ana Z. Soto, Laura Corrigan, David W. Sims, Martin J. Genner, and Stefano Mariani. 2019. “Non-Specific Amplification Compromises Environmental DNA Metabarcoding with COI.” Methods in Ecology and Evolution 10(11):1985–2001. doi: 10.1111/2041-210X.13276.

Costa, Filipe O., and Gary R. Carvalho. 2007. “The Barcode of Life Initiative: Synopsis and Prospective Societal Impacts of DNA Barcoding of Fish.” *Genomics*, Society and Policy 3(2):29. doi: 10.1186/1746-5354-3-2-29.

Davidson, Nick C. 2014. “How Much Wetland Has the World Lost? Long-Term and Recent Trends in Global Wetland Area.” Marine and Freshwater Research 65(10):934–41. doi: 10.1071/MF14173.

Deagle, Bruce E., Simon N. Jarman, Eric Coissac, François Pompanon, and Pierre Taberlet. 2014. “DNA Metabarcoding and the Cytochrome c Oxidase Subunit I Marker: Not a Perfect Match.” Biology Letters 10(9):20140562. doi: 10.1098/rsbl.2014.0562.

Deiner, Kristy, and Florian Altermatt. 2014. “Transport Distance of Invertebrate Environmental DNA in a Natural River.” PLOS ONE 9(2):e88786. doi: 10.1371/journal.pone.0088786.

Deiner, Kristy, Emanuel A. Fronhofer, Elvira Mächler, Jean-Claude Walser, and Florian Altermatt. 2016. “Environmental DNA Reveals That Rivers Are Conveyer Belts of Biodiversity Information.” Nature Communications 7(1):12544. doi: 10.1038/ncomms12544.

Dejean, Tony, Alice Valentini, Antoine Duparc, Stéphanie Pellier-Cuit, François Pompanon, Pierre Taberlet, and Claude Miaud. 2011. “Persistence of Environmental DNA in Freshwater Ecosystems.” PLOS ONE 6(8):e23398. doi: 10.1371/journal.pone.0023398.

Duarte, Sofia, Barbara R. Leite, Maria João Feio, Filipe O. Costa, and Ana Filipa Filipe. 2021. “Integration of DNA-Based Approaches in Aquatic Ecological Assessment Using Benthic Macroinvertebrates.” Water 13(3):331. doi: 10.3390/w13030331.

Dudgeon, David, Angela H. Arthington, Mark O. Gessner, Zen-Ichiro Kawabata, Duncan J. Knowler, Christian Lévêque, Robert J. Naiman, Anne-Hélène Prieur-Richard, Doris Soto, Melanie L. J. Stiassny, and Caroline A. Sullivan. 2006. “Freshwater Biodiversity: Importance, Threats, Status and Conservation Challenges.” Biological Reviews 81(2):163–82. doi: 10.1017/S1464793105006950.

Elbrecht, Vasco, Thomas W. A. Braukmann, Natalia V. Ivanova, Sean W. J. Prosser, Mehrdad Hajibabaei, Michael Wright, Evgeny V. Zakharov, Paul D. N. Hebert, and Dirk Steinke. 2019. “Validation of COI Metabarcoding Primers for Terrestrial Arthropods.” PeerJ 7:e7745. doi: 10.7717/peerj.7745.

Elbrecht, Vasco, and Florian Leese. 2016. “PrimerMiner: An r Package for Development and in Silico Validation of DNA Metabarcoding Primers.” Methods in Ecology and Evolution 8(5):622–26. doi: 10.1111/2041-210X.12687.

Elbrecht, Vasco, and Florian Leese. 2017. “Validation and Development of COI Metabarcoding Primers for Freshwater Macroinvertebrate Bioassessment.” Frontiers in Environmental Science 5. doi: 10.3389/fenvs.2017.00011.

Estensmo, Eva Lena F., Sundy Maurice, Luis Morgado, Pedro M. Martin-Sanchez, Inger Skrede, and Håvard Kauserud. 2021. “The Influence of Intraspecific Sequence Variation during DNA Metabarcoding: A Case Study of Eleven Fungal Species.” Molecular Ecology Resources 21(4):1141–48. doi: 10.1111/1755-0998.13329.

Fediajevaite, Julija, Victoria Priestley, Richard Arnold, and Vincent Savolainen. 2021. “Metalanalysis Shows That Environmental DNA Outperforms Traditional Surveys, but Warrants Better Reporting Standards.” Ecology and Evolution 11(9):4803–15. doi: 10.1002/ece3.7382.

Ficetola, Gentile F., Johan Pansu, Aurélie Bonin, Eric Coissac, Charline GiguetlCovex, Marta De Barba, Ludovic Gielly, Carla M. Lopes, Frédéric Boyer, François Pompanon, Gilles Rayé, and Pierre Taberlet. 2015. “Replication Levels, False Presences and the Estimation of the Presence/Absence from eDNA Metabarcoding Data.” Molecular Ecology Resources 15(3):543–56. doi: 10.1111/1755-0998.12338.

Ficetola, Gentile Francesco, Eric Coissac, Stéphanie Zundel, Tiayyba Riaz, Wasim Shehzad, Julien Bessière, Pierre Taberlet, and François Pompanon. 2010. “An In Silico Approach for the Evaluation of DNA Barcodes.” BMC Genomics 11(1):434. doi: 10.1186/1471-2164-11-434.

Ficetola, Gentile Francesco, and Pierre Taberlet. 2022. *Towards All-Inclusive Community Ecology via DNA Metabarcoding*. *preprint*. Preprints. doi: 10.22541/au.166546664.45118988/v1.

Folmer, O., W. R. Hoeh, M. B. Black, and R. C. Vrijenhoek. 1994. “Conserved Primers for PCR Amplification of Mitochondrial DNA from Different Invertebrate Phyla.” Molecular Marine Biology and Biotechnology 3:294–99.

Freeland, Joanna R. 2017. “The Importance of Molecular Markers and Primer Design When Characterizing Biodiversity from Environmental DNA.” Genome 60(4):358–74. doi: 10.1139/gen-2016-0100.

Galan, Maxime, Maria Razzauti, Emilie Bard, Maria Bernard, Carine Brouat, Nathalie Charbonnel, Alexandre Dehne-Garcia, Anne Loiseau, Caroline Tatard, Lucie Tamisier, Muriel Vayssier-Taussat, Helene Vignes, and Jean-François Cosson. 2016. “16S rRNA Amplicon Sequencing for Epidemiological Surveys of Bacteria in Wildlife.” mSystems 1(4):e00032–16. doi: 10.1128/mSystems.00032-16.

Gibson, Thomas I., Gary Carvalho, Amy Ellison, Enrica Gargiulo, Tristan Hatton-Ellis, Lori Lawson-Handley, Stefano Mariani, Rupert A. Collins, Graham Sellers, Marco Antonio Distaso, Carlo Zampieri, and Simon Creer. 2023. “Environmental DNA Metabarcoding for Fish Diversity Assessment in a Macrotidal Estuary: A Comparison with Established Fish Survey Methods.” *Estuarine*, Coastal and Shelf Science 294:108522. doi: 10.1016/j.ecss.2023.108522.

Goutte, Aurélie, Noëlie Molbert, Sabrina Guérin, Robin Richoux, and Vincent Rocher. 2020. “Monitoring Freshwater Fish Communities in Large Rivers Using Environmental DNA Metabarcoding and a Long-Term Electrofishing Survey.” Journal of Fish Biology 97(2):444–52. doi: 10.1111/jfb.14383.

Greenhalgh, Jack A., Rupert A. Collins, Duncan E. Edgley, Martin J. Genner, Jan Hindle, Gareth Jones, Lesley Loughlin, Maire O’donnel, Michael J. Sweet, and Richard W. Battarbee. 2022. “Environmental DNA-Based Methods Detect the Invasion Front of an Advancing Signal Crayfish Population.” Environmental DNA 4(3):596–607. doi: 10.1002/edn3.280.

Guillera-Arroita, Gurutzeta, José Joaquín Lahoz-Monfort, Anthony R. van Rooyen, Andrew R. Weeks, and Reid Tingley. 2017. “Dealing with False-Positive and False-Negative Errors about Species Occurrence at Multiple Levels.” Methods in Ecology and Evolution 8(9):1081–91. doi: 10.1111/2041-210X.12743.

Hallam, Jane, Elizabeth L. Clare, John Iwan Jones, and Julia J. Day. 2021. “Biodiversity Assessment across a Dynamic Riverine System: A Comparison of eDNA Metabarcoding versus Traditional Fish Surveying Methods.” Environmental DNA 3(6):1247–66. doi: 10.1002/edn3.241.

Hebert, Paul D. N., Sujeevan Ratnasingham, and Jeremy R. de Waard. 2003. “Barcoding Animal Life: Cytochrome c Oxidase Subunit 1 Divergences among Closely Related Species.” Proceedings of the Royal Society of London. Series B: Biological Sciences 270(suppl_1):S96–99. doi: 10.1098/rsbl.2003.0025.

Higgins, S. N., and M. J. Vander Zanden. 2010. “What a Difference a Species Makes: A Meta– Analysis of Dreissenid Mussel Impacts on Freshwater Ecosystems.” Ecological Monographs 80(2):179–96. doi: 10.1890/09-1249.1.

Hobern, Donald. 2021. “BIOSCAN: DNA Barcoding to Accelerate Taxonomy and Biogeography for Conservation and Sustainability.” Genome 64(3):161–64. doi: 10.1139/gen-2020-0009.

Iwasaki, Wataru, Tsukasa Fukunaga, Ryota Isagozawa, Koichiro Yamada, Yasunobu Maeda, Takashi P. Satoh, Tetsuya Sado, Kohji Mabuchi, Hirohiko Takeshima, Masaki Miya, and Mutsumi Nishida. 2013. “MitoFish and MitoAnnotator: A Mitochondrial Genome Database of Fish with an Accurate and Automatic Annotation Pipeline.” Molecular Biology and Evolution 30(11):2531–40. doi: 10.1093/molbev/mst141.

Ji, Fenfen, Dingyi Han, Liang Yan, Saihong Yan, Jinmiao Zha, and Jianzhong Shen. 2022. “Assessment of Benthic Invertebrate Diversity and River Ecological Status along an Urbanized Gradient Using Environmental DNA Metabarcoding and a Traditional Survey Method.” Science of The Total Environment 806:150587. doi: 10.1016/j.scitotenv.2021.150587.

Karatayev, Alexander Y., and Lyubov E. Burlakova. 2022. “Dreissena in the Great Lakes: What Have We Learned in 30 Years of Invasion.” Hydrobiologia. doi: 10.1007/s10750-022-04990-x.

Kearse, Matthew, Richard Moir, Amy Wilson, Steven Stones-Havas, Matthew Cheung, Shane Sturrock, Simon Buxton, Alex Cooper, Sidney Markowitz, Chris Duran, Tobias Thierer, Bruce Ashton, Peter Meintjes, and Alexei Drummond. 2012. “Geneious Basic: An Integrated and Extendable Desktop Software Platform for the Organization and Analysis of Sequence Data.” Bioinformatics 28(12):1647–49. doi: 10.1093/bioinformatics/bts199.

Keck, François, Rosetta C. Blackman, Raphael Bossart, Jeanine Brantschen, Marjorie Couton, Samuel Hürlemann, Dominik Kirschner, Nadine Locher, Heng Zhang, and Florian Altermatt. 2021. Meta-Analysis Shows Both Congruence and Complementarity of DNA Metabarcoding to Traditional Methods for Biological Community Assessment. doi: 10.1101/2021.06.29.450286.

Keck, François, Marjorie Couton, and Florian Altermatt. 2023. “Navigating the Seven Challenges of Taxonomic Reference Databases in Metabarcoding Analyses.” Molecular Ecology Resources 23(4):742–55. doi: 10.1111/1755-0998.13746.

Keller, Abigail G., Emily W. Grason, P. Sean McDonald, Ana Ramón-Laca, and Ryan P. Kelly. 2022. “Tracking an Invasion Front with Environmental DNA.” Ecological Applications 32(4):e2561. doi: 10.1002/eap.2561.

Kelly, Ryan P., Andrew Olaf Shelton, and Ramón Gallego. 2019. “Understanding PCR Processes to Draw Meaningful Conclusions from Environmental DNA Studies.” Scientific Reports 9(1):12133. doi: 10.1038/s41598-019-48546-x.

Klymus, Katy E., Nathaniel T. Marshall, and Carol A. Stepien. 2017. “Environmental DNA (eDNA) Metabarcoding Assays to Detect Invasive Invertebrate Species in the Great Lakes.” PLoS ONE 12(5):e0177643. doi: 10.1371/journal.pone.0177643.

Klymus, Katy E., Catherine A. Richter, Nathan Thompson, and Jo Ellen Hinck. 2017. “Metabarcoding of Environmental DNA Samples to Explore the Use of Uranium Mine Containment Ponds as a Water Source for Wildlife.” Diversity 9(4):54. doi: 10.3390/d9040054.

Kozich, James J., Sarah L. Westcott, Nielson T. Baxter, Sarah K. Highlander, and Patrick D. Schloss. 2013. “Development of a Dual-Index Sequencing Strategy and Curation Pipeline for Analyzing Amplicon Sequence Data on the MiSeq Illumina Sequencing Platform.” Applied and Environmental Microbiology AEM.01043–13. doi: 10.1128/AEM.01043-13.

Krehenwinkel, Henrik, Madeline Wolf, Jun Ying Lim, Andrew J. Rominger, Warren B. Simison, and Rosemary G. Gillespie. 2017. “Estimating and Mitigating Amplification Bias in Qualitative and Quantitative Arthropod Metabarcoding.” Scientific Reports 7(1):17668. doi: 10.1038/s41598-017-17333-x.

Laporte, Martin, Bérénice Bougas, Guillaume Côté, Olivier Champoux, Yves Paradis, Jean Morin, and Louis Bernatchez. 2020. “Caged Fish Experiment and Hydrodynamic Bidimensional Modeling Highlight the Importance to Consider 2D Dispersion in Fluvial Environmental DNA Studies.” Environmental DNA 2(3):362–72. doi: 10.1002/edn3.88.

Leese, Florian, Mandy Sander, Dominik Buchner, Vasco Elbrecht, Peter Haase, and Vera M. A. Zizka. 2021. “Improved Freshwater Macroinvertebrate Detection from Environmental DNA through Minimized Nontarget Amplification.” Environmental DNA 3(1):261–76. doi: 10.1002/edn3.177.

Lenth, R. 2023. “Emmeans: Estimated Marginal Means, Aka Least-Squares Means. R Package.”

Leray, Matthieu, Joy Y. Yang, Christopher P. Meyer, Suzanne C. Mills, Natalia Agudelo, Vincent Ranwez, Joel T. Boehm, and Ryuji J. Machida. 2013. “A New Versatile Primer Set Targeting a Short Fragment of the Mitochondrial COI Region for Metabarcoding Metazoan Diversity: Application for Characterizing Coral Reef Fish Gut Contents.” Frontiers in Zoology 10(1):34. doi: 10.1186/1742-9994-10-34.

Lopez, Mark Louie D., Matthew Bonderud, Michael J. Allison, Findlay MacDermid, Erin J. Ussery, Mark E. McMaster, Ave Dersch, Kasia J. Staniszewska, Colin A. Cooke, Paul Drevnick, and Caren C. Helbing. 2023. “qPCR-Based eDNA Workflow for Humic-Rich Lake Sediments: Combined Use of Sedimentary DNA (sedDNA) and Indigenous Knowledge in Reconstructing Historical Fish Records.” Ecological Indicators 155:111014. doi: 10.1016/j.ecolind.2023.111014.

Lyet, Arnaud, Loïc Pellissier, Alice Valentini, Tony Dejean, Abigail Hehmeyer, and Robin Naidoo. 2021. “eDNA Sampled from Stream Networks Correlates with Camera Trap Detection Rates of Terrestrial Mammals.” Scientific Reports 11(1):11362. doi: 10.1038/s41598-021-90598-5.

Macher, Till-Hendrik, Robin Schütz, Atakan Yildiz, Arne J. Beermann, and Florian Leese. 2023. “Evaluating Five Primer Pairs for Environmental DNA Metabarcoding of Central European Fish Species Based on Mock Communities.” Metabarcoding and Metagenomics 7:e103856. doi: 10.3897/mbmg.7.103856.

Magoč, Tanja, and Steven L. Salzberg. 2011. “FLASH: Fast Length Adjustment of Short Reads to Improve Genome Assemblies.” Bioinformatics 27(21):2957–63. doi: 10.1093/bioinformatics/btr507.

Marbek. 2023. Assessing the Economic Value of Protecting the Great Lakes Ecosystems. Ontario Ministry of Environment.

Marinchel, Nadia, Alexis Marchesini, Davide Nardi, Matteo Girardi, Silvia Casabianca, Cristiano Vernesi, and Antonella Penna. 2023. “Mock Community Experiments Can Inform on the Reliability of eDNA Metabarcoding Data: A Case Study on Marine Phytoplankton.” Scientific Reports 13(1):20164. doi: 10.1038/s41598-023-47462-5.

Marques, Virginie, Tristan Milhau, Camille Albouy, Tony Dejean, Stéphanie Manel, David Mouillot, and Jean-Baptiste Juhel. 2021. “GAPeDNA: Assessing and Mapping Global Species Gaps in Genetic Databases for eDNA Metabarcoding.” Diversity and Distributions 27(10):1880–92. doi: 10.1111/ddi.13142.

Mathon, Laetitia, Alice Valentini, Pierre-Edouard Guérin, Eric Normandeau, Cyril Noel, Clément Lionnet, Emilie Boulanger, Wilfried Thuiller, Louis Bernatchez, David Mouillot, Tony Dejean, and Stéphanie Manel. 2021. “Benchmarking Bioinformatic Tools for Fast and Accurate eDNA Metabarcoding Species Identification.” Molecular Ecology Resources 21(7):2565–79. doi: 10.1111/1755-0998.13430.

McManus, George B., and Laura A. Katz. 2009. “Molecular and Morphological Methods for Identifying Plankton: What Makes a Successful Marriage?” Journal of Plankton Research 31(10):1119–29. doi: 10.1093/plankt/fbp061.

McMurdie, Paul J., and Susan Holmes. 2014. “Waste Not, Want Not: Why Rarefying Microbiome Data Is Inadmissible.” PLOS Computational Biology 10(4):e1003531. doi: 10.1371/journal.pcbi.1003531.

Meiklejohn, Kelly A., Natalie Damaso, and James M. Robertson. 2019. “Assessment of BOLD and GenBank – Their Accuracy and Reliability for the Identification of Biological Materials.” PLoS ONE 14(6):e0217084. doi: 10.1371/journal.pone.0217084.

Meusnier, Isabelle, Gregory AC Singer, Jean-François Landry, Donal A. Hickey, Paul DN Hebert, and Mehrdad Hajibabaei. 2008. “A Universal DNA Mini-Barcode for Biodiversity Analysis.” BMC Genomics 9(1):214. doi: 10.1186/1471-2164-9-214.

Miya, M., Y. Sato, T. Fukunaga, T. Sado, J. Y. Poulsen, K. Sato, T. Minamoto, S. Yamamoto, H. Yamanaka, H. Araki, M. Kondoh, and W. Iwasaki. 2015. “MiFish, a Set of Universal PCR Primers for Metabarcoding Environmental DNA from Fishes: Detection of More than 230 Subtropical Marine Species.” Royal Society Open Science 2(7):150088. doi: 10.1098/rsos.150088.

Miya, Masaki, Ryo O. Gotoh, and Tetsuya Sado. 2020. “MiFish Metabarcoding: A High-Throughput Approach for Simultaneous Detection of Multiple Fish Species from Environmental DNA and Other Samples.” Fisheries Science 86(6):939–70. doi: 10.1007/s12562-020-01461-x.

Piczak, Morgan L., Denielle Perry, Steven J. Cooke, Ian Harrison, Silvia Benitez, Aaron Koning, Li Peng, Peter Limbu, Karen E. Smokorowski, Sergio Salinas-Rodriguez, John D. Koehn, and Irena F. Creed. 2023. “Protecting and Restoring Habitats to Benefit Freshwater Biodiversity.” Environmental Reviews. doi: 10.1139/er-2023-0034.

Porter, Teresita M., and Mehrdad Hajibabaei. 2018. “Over 2.5 Million COI Sequences in GenBank and Growing.” PLoS ONE 13(9):e0200177. doi: 10.1371/journal.pone.0200177.

R Core Team. 2022. “R: A Language and Environment for Statistical Computing.”

Ratnasingham, Sujeevan, and P. D. N. Hebert. 2007. “Bold: The Barcode of Life Data System (Http://Www.Barcodinglife.Org).” Molecular Ecology Notes 7(3):355–64. doi: 10.1111/j.1471-8286.2007.01678.x.

Reid, Andrea J., Andrew K. Carlson, Irena F. Creed, Erika J. Eliason, Peter A. Gell, Pieter T. J. Johnson, Karen A. Kidd, Tyson J. MacCormack, Julian D. Olden, Steve J. Ormerod, John P. Smol, William W. Taylor, Klement Tockner, Jesse C. Vermaire, David Dudgeon, and Steven J. Cooke. 2019. “Emerging Threats and Persistent Conservation Challenges for Freshwater Biodiversity.” Biological Reviews 94(3):849–73. doi: 10.1111/brv.12480.

Ricciardi, A., F. G. Whoriskey, and J. B. Rasmussen. 1996. “Impact of the (*Dreissena*) Invasion on Native Unionid Bivalves in the Upper St. Lawrence River.” Canadian Journal of Fisheries and Aquatic Sciences 53(6):1434–44. doi: 10.1139/f96-068.

Rognes, Torbjørn, Tomáš Flouri, Ben Nichols, Christopher Quince, and Frédéric Mahé. 2016. “VSEARCH: A Versatile Open Source Tool for Metagenomics.” PeerJ 4:e2584. doi: 10.7717/peerj.2584.

Roy, Mélanie, Valérie Belliveau, Nicholas E. Mandrak, and Nellie Gagné. 2018. “Development of Environmental DNA (eDNA) Methods for Detecting High-Risk Freshwater Fishes in Live Trade in Canada.” Biological Invasions 20(2):299–314. doi: 10.1007/s10530-017-1532-z.

Sales, Naiara Guimarães, Maisie B. McKenzie, Joseph Drake, Lynsey R. Harper, Samuel S. Browett, Ilaria Coscia, Owen S. Wangensteen, Charles Baillie, Emma Bryce, Deborah A. Dawson, Erinma Ochu, Bernd Hänfling, Lori Lawson Handley, Stefano Mariani, Xavier Lambin, Christopher Sutherland, and Allan D. McDevitt. 2020. “Fishing for Mammals: Landscape-Level Monitoring of Terrestrial and Semi-Aquatic Communities Using eDNA from Riverine Systems.” Journal of Applied Ecology 57(4):707–16. doi: 10.1111/1365-2664.13592.

Sato, Yukuto, Masaki Miya, Tsukasa Fukunaga, Tetsuya Sado, and Wataru Iwasaki. 2018. “MitoFish and MiFish Pipeline: A Mitochondrial Genome Database of Fish with an Analysis Pipeline for Environmental DNA Metabarcoding.” Molecular Biology and Evolution 35(6):1553–55. doi: 10.1093/molbev/msy074.

Shirazi, Sabrina, Rachel S. Meyer, and Beth Shapiro. 2021. “Revisiting the Effect of PCR Replication and Sequencing Depth on Biodiversity Metrics in Environmental DNA Metabarcoding.” Ecology and Evolution 11(22):15766–79. doi: 10.1002/ece3.8239.

Smith, Dylan P., and Kabir G. Peay. 2014. “Sequence Depth, Not PCR Replication, Improves Ecological Inference from Next Generation DNA Sequencing.” PLoS ONE 9(2):e90234. doi: 10.1371/journal.pone.0090234.

Smith, Sigrid D. P., David B. Bunnell, G. A. Burton, Jan J. H. Ciborowski, Alisha D. Davidson, Caitlin E. Dickinson, Lauren A. Eaton, Peter C. Esselman, Mary Anne Evans, Donna R. Kashian, Nathan F. Manning, Peter B. McIntyre, Thomas F. Nalepa, Alicia Pérez-Fuentetaja, Alan D. Steinman, Donald G. Uzarski, and J. David Allan. 2019. “Evidence for Interactions among Environmental Stressors in the Laurentian Great Lakes.” Ecological Indicators 101:203–11. doi: 10.1016/j.ecolind.2019.01.010.

Stadhouders, Ralph, Suzan D. Pas, Jeer Anber, Jolanda Voermans, Ted H. M. Mes, and Martin Schutten. 2010. “The Effect of Primer-Template Mismatches on the Detection and Quantification of Nucleic Acids Using the 5′ Nuclease Assay.” The Journal of Molecular Diagnostics 12(1):109–17. doi: 10.2353/jmoldx.2010.090035.

Stat, Michael, Megan J. Huggett, Rachele Bernasconi, Joseph D. DiBattista, Tina E. Berry, Stephen J. Newman, Euan S. Harvey, and Michael Bunce. 2017. “Ecosystem Biomonitoring with eDNA: Metabarcoding across the Tree of Life in a Tropical Marine Environment.” Scientific Reports 7(1):12240. doi: 10.1038/s41598-017-12501-5.

Thalinger, Bettina, Kristy Deiner, Lynsey R. Harper, Helen C. Rees, Rosetta C. Blackman, Daniela Sint, Michael Traugott, Caren S. Goldberg, and Kat Bruce. 2021. “A Validation Scale to Determine the Readiness of Environmental DNA Assays for Routine Species Monitoring.” Environmental DNA 3(4):823–36. doi: 10.1002/edn3.189.

Tournayre, Orianne, Maxime Leuchtmann, Ondine FilippilCodaccioni, Marine Trillat, Sylvain Piry, Dominique Pontier, Nathalie Charbonnel, and Maxime Galan. 2020. “In Silico and Empirical Evaluation of Twelve Metabarcoding Primer Sets for Insectivorous Diet Analyses.” Ecology and Evolution 10(13):6310–32. doi: 10.1002/ece3.6362.

Turner, Cameron R., Derryl J. Miller, Kathryn J. Coyne, and Joel Corush. 2014. “Improved Methods for Capture, Extraction, and Quantitative Assay of Environmental DNA from Asian Bigheaded Carp (*Hypophthalmichthys Spp.*).” PLOS ONE 9(12):e114329. doi: 10.1371/journal.pone.0114329.

Tzafesta, Eftychia, Benedetta Saccomanno, Francesco Zangaro, Maria Rosaria Vadrucci, Valeria Specchia, and Maurizio Pinna. 2022. “DNA Barcode Gap Analysis for Multiple Marker Genes for Phytoplankton Species Biodiversity in Mediterranean Aquatic Ecosystems.” Biology 11(9):1277. doi: 10.3390/biology11091277.

Valentini, Alice, Pierre Taberlet, Claude Miaud, Raphaël Civade, Jelger Herder, Philip Francis Thomsen, Eva Bellemain, Aurélien Besnard, Eric Coissac, Frédéric Boyer, Coline Gaboriaud, Pauline Jean, Nicolas Poulet, Nicolas Roset, Gordon H. Copp, Philippe Geniez, Didier Pont, Christine Argillier, Jean-Marc Baudoin, Tiphaine Peroux, Alain J. Crivelli, Anthony Olivier, Manon Acqueberge, Matthieu Le Brun, Peter R. Møller, Eske Willerslev, and Tony Dejean. 2016. “Next-Generation Monitoring of Aquatic Biodiversity Using Environmental DNA Metabarcoding.” Molecular Ecology 25(4):929–42. doi: 10.1111/mec.13428.

Wacker, Sebastian, Frode Fossøy, Bjørn Mejdell Larsen, Hege Brandsegg, Rolf Sivertsgård, and Sten Karlsson. 2019. “Downstream Transport and Seasonal Variation in Freshwater Pearl Mussel (*Margaritifera Margaritifera*) eDNA Concentration.” Environmental DNA 1(1):64–73. doi: 10.1002/edn3.10.

Weigand, Hannah, Arne J. Beermann, Fedor Čiampor, Filipe O. Costa, Zoltán Csabai, Sofia Duarte, Matthias F. Geiger, Michał Grabowski, Frédéric Rimet, Björn Rulik, Malin Strand, Nikolaus Szucsich, Alexander M. Weigand, Endre Willassen, Sofia A. Wyler, Agnès Bouchez, Angel Borja, Zuzana Čiamporová-Zaťovičová, Sónia Ferreira, Klaas-Douwe B. Dijkstra, Ursula Eisendle, Jörg Freyhof, Piotr Gadawski, Wolfram Graf, Arne Haegerbaeumer, Berry B. van der Hoorn, Bella Japoshvili, Lujza Keresztes, Emre Keskin, Florian Leese, Jan N. Macher, Tomasz Mamos, Guy Paz, Vladimir Pešić, Daniela Maric Pfannkuchen, Martin Andreas Pfannkuchen, Benjamin W. Price, Buki Rinkevich, Marcos A. L. Teixeira, Gábor Várbíró, and Torbjørn Ekrem. 2019. “DNA Barcode Reference Libraries for the Monitoring of Aquatic Biota in Europe: Gap-Analysis and Recommendations for Future Work.” Science of The Total Environment 678:499–524. doi: 10.1016/j.scitotenv.2019.04.247.

Yu, Douglas W., Yinqiu Ji, Brent C. Emerson, Xiaoyang Wang, Chengxi Ye, Chunyan Yang, and Zhaoli Ding. 2012. “Biodiversity Soup: Metabarcoding of Arthropods for Rapid Biodiversity Assessment and Biomonitoring.” Methods in Ecology and Evolution 4:613–23. doi: 10.1111/j.2041-210X.2012.00198.x@10.1111/(ISSN)1365-2435.ECOLOGYCHINA.

Zaiko, Anastasija, Xavier Pochon, Eva Garcia-Vazquez, Sergej Olenin, and Susanna A. Wood. 2018. “Advantages and Limitations of Environmental DNA/RNA Tools for Marine Biosecurity: Management and Surveillance of Non-Indigenous Species.” Frontiers in Marine Science 5.

Zhan, Aibin, Martin Hulák, Francisco Sylvester, Xiaoting Huang, Abisola A. Adebayo, Cathryn L. Abbott, Sarah J. Adamowicz, Daniel D. Heath, Melania E. Cristescu, and Hugh J. MacIsaac. 2013. “High Sensitivity of 454 Pyrosequencing for Detection of Rare Species in Aquatic Communities.” Methods in Ecology and Evolution 4(6):558–65. doi: 10.1111/2041-210X.12037.

